# Dual thalamic drive defines the intra-amygdala wiring complexity

**DOI:** 10.64898/2026.06.29.735253

**Authors:** Ákos Babiczky, Péter Berki, Liu Mengxing, Kinga Kocsis, Aletta Magyar, Anna V. Bakacsi, Boglárka Barsy, Judit Berczik, Sándor Borbély, Antal Berényi, Francisco Clascá, Pedro M. Paz-Alonso, Ferenc Mátyás

**Author notes:** Corresponding author: Ferenc Mátyás.

## Abstract

The amygdala plays a key role in affective behaviors by integrating incoming signals and conveying them towards subcortical output regions. Current models suggest a serial information flow within the amygdala from the lateral and basolateral towards the central subnuclei driven by incoming thalamic and cortical excitation. However, due to the lack of universally accepted parcellation principles, the precise connectivity and thus, the signal propagation within the circuit remain debated. Using a molecular-based parcellation, our subnucleus-specific anatomical and electrophysiological mapping revealed a previously overlooked complexity in the intra-amygdala wiring pattern in mice. Specifically, the intra-amygdala signal transfer relies on separate lateral thalamus-driven routes via the lateral subnucleus mostly bypassing the anterior basolateral and centrolateral subnuclei. In contrast, the anterior basolateral nucleus, innervated by the dorsal midline thalamus, supplies mostly extra-, but not intra-amygdala routes. We also demonstrated that a similar dual thalamo-cortico-amygdala organization exists in the human brain. Collectively, our findings identified unconventional amygdala wiring principles challenging the traditional serial lateral-basolateral-central stream model which can redefine our understanding of behaviorally relevant intra-amygdala computations.

## Introduction

Affective processes promote survival, decision-making and social interaction, placing them in the centre of neuroscientific attention for decades^1–3^. The amygdala, a multidomain brain region became widely known as a central hub for regulating affective behaviors by integrating various thalamic and cortical information and providing efferent signals to subcortical areas directly involved in executive functions^4–7^. Direct thalamo-amygdala connections are often implicated in simply relaying relatively unprocessed sensory information^8–10^. However, recent studies have revealed the involvement of functionally different amygdala-projecting thalamic areas, such as the dorsal (dMT) and ventral midline (vMT), gustatory (GusT) and lateral thalamic nuclei (LT), in more complex affective information processing^11–20^. Although most of these nuclei were reported to innervate several nuclei in the amygdala^16,19–23^, the precise subnucleus-selective description of thalamo-amygdala connections, and so, our understanding of intra-amygdala signal processing is still incomplete.

The amygdala is often depicted as a single processing unit with a rather serial internal information flow, despite its known compartmentalized nature and complex wiring dynamics^6,9,10,24–28^. In brief, it is suggested that incoming signals first reach the cortical-like amygdala nuclei, such as the lateral (LA) and the anterior basolateral nucleus (BLA), which, in turn, innervate the striatal-like central amygdala (CeA). Then, CeA directly governs behavioral, autonomic and hormonal responses through its efferent connections targeting various subcortical regions. However, both the cortical- and striatal-like amygdala can be further divided into several subnuclei possessing markedly different anatomical and functional characteristics, although a universally accepted and used parcellation approach is lacking. Without such a unified framework, it is challenging to compare results and to draw conclusions from experimental data concerning the connectivity of subnuclei, and in turn, the flow of information processing within the amygdala complex. Inconsistencies in the definition of individual amygdala subnuclei might be a reason behind seemingly contradictory functional findings about certain intra-amygdala pathways, such as the one between the basolateral and central nuclei^29–33^.

Taken together, thalamic afferents in the amygdala are often treated as a single entity, despite their clear functional and anatomical differences. Furthermore, they are suggested to ultimately drive a serial information processing pathway within the amygdala (LA→BLA→CeA route), although individual amygdala subnuclei were linked to substantially different functions. Such heterogeneity implies the existence of a more complex thalamo-amygdala network and intra-amygdala wiring than what is currently proposed.

Using a combination of classical and intersectional anatomical tracing techniques as well as high-density electrophysiological recordings in mice, here we report a previously overlooked complexity within the amygdala, defined by the two major thalamic afferents. By introducing a novel, reproducible amygdala parcellation approach based on endogenous protein expression, we demonstrate that LT drives various sensory-related intra-amygdala pathways via the LA, while the dMT and the BLA provide a link towards a brain-wide limbic network outside the amygdala. This duality, which we also describe in the human brain, would enable parallel processing of emotionally relevant information through the thalamo-amygdala network.

## Results

### Duality in the thalamo-cortico-amygdala organization is evolutionarily conserved

To systematically map the sources of thalamic and cortical inputs to the amygdala, we injected large deposits of classical retrograde tracer cholera toxin B (CTB) covering most of the amygdala in mice (Fig. 1a). In the anterior part of the thalamus, we found retrogradely labelled cells in the dorsal midline region (dMT), most prominently in the paraventricular thalamic nucleus (PVT). Caudally, the majority of labelled cells were found in two nuclei, the suprageniculate (SG) and the posterior intralaminar nucleus (PIL) (Fig. 1b and Extended Data Fig. 1). For simplicity, we will refer to these nuclei collectively as lateral thalamus (LT)^16^. At the level of the neocortex, we found amygdala-projecting neurons in the medial part of the prefrontal cortex (PFC), including the anterior cingulate (Cg1-2), prelimbic (PrL) and infralimbic (IL) regions, and in higher-order regions of the temporal cortex (TeCx), most densely in the temporal association cortex (TeA) (Fig. 1c and Extended Data Fig. 1). As these cortical regions are also connected to dMT or LT^34–36^, we hypothesized a duality in the organization of the thalamo-cortico-amygdala network.

**Figure 1:**
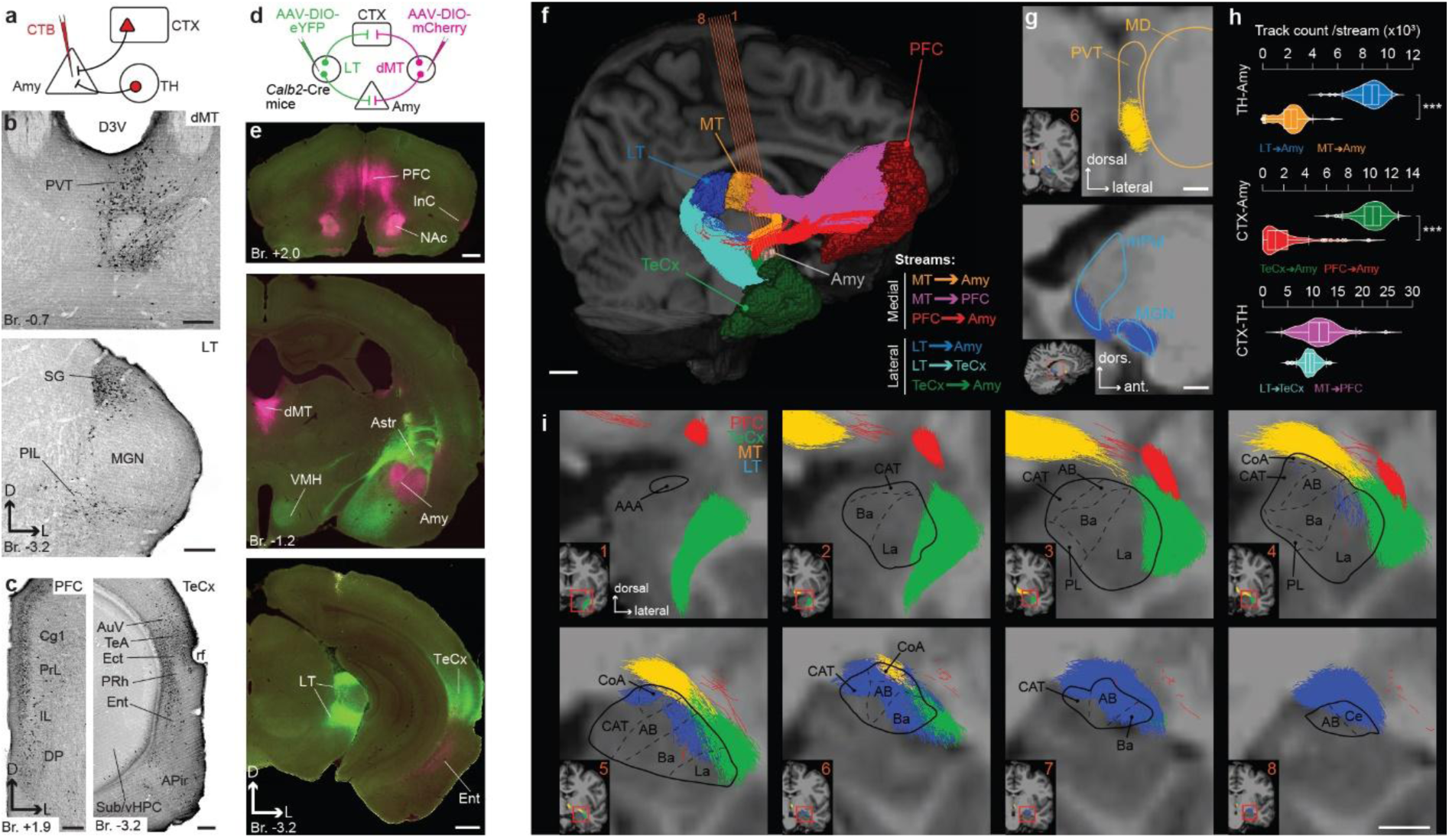
Duality in the thalamo-cortico-amygdala organization is evolutionarily conserved. **a,** Experimental design for retrograde (cholera toxin B; CTB) tracings in the thalamo-cortico-amygdala network in mice (*n* = 3). **b**,**c**, Representative brightfield images showing the distribution of amygdala-projecting thalamic (**b**) and cortical (**c**) neurons. Scale bars, 200 µm. **d,** Experimental design for AAV-mediated anterograde tracings to simultaneously label the lateral (LT) and the dorsomedial thalamus (dMT)-linked circuits in mice (*n* = 4). **e,** Distribution of the two major thalamic afferents in the cortical regions and the amygdala: LT (green) and dMT (magenta). Scale bars, 500 µm. **f,** 3D streamline visualizations of human brain tractography showing lateral (cold colors) and medial (warm colors) thalamo-amygdala, cortico-amygdala and thalamo-cortical streams. Orange lines (1-8) indicate coronal sections for (g,i). Scale bar, 1 cm. **g,** Representative ROIs of the MT and the LT regions in human. Insets: positions at the macroscopic level. Scale bars, 5 mm. **h,** Violin plots showing the quantification of tract counts in each stream (*n* = 113 subjects; Kruskal-Wallis test, *H(5)* = 514.1, *P* = 7.283*10^-109^, Dunn’s post-hoc test, \*\*\**P* < 0.001). **i,** Magnified streamline visualization of PFC- (red), TeCx- (green), MT- (orange) and LT-amygdala (blue) tractographies in coronal sections at 8 different AP levels. Insets: position at the macroscopic level. Whole amygdala (solid line) and amygdala subnuclei borders (dashed lines) are highlighted. Scale bars, 5 mm. All human images in this figure are taken from one representative subject. For detailed statistical data, abbreviations and raw data see *Supplementary Tables* and *Source Data Tables*.

As majority of thalamic innervation to the amygdala originates in calretinin (Calr)-expressing neurons (Extended Data Fig. 2)^16,23^, we selectively labelled LT and dMT efferents using Cre-dependent AAV vectors carrying either eYFP or mCherry in Calb2-Cre mice (Fig. 1d). Confirming our hypothesis, while LT axons mostly innervated the TeCx, dMT axons appeared predominantly in the PFC, and not vice versa. On the other hand, both thalamic regions innervated the amygdala, although their axonal distributions showed clear differences (Fig. 1e and Extended Data Fig. 3). Collectively, our anatomical investigations in mice showed the existence of a lateral (LT-TeCx-Amy) and a medial (dMT-PFC-Amy) stream in the thalamo-cortico-amygdala circuitry, originating in Calr-positive thalamic neurons. These streams resembled the traditional “low road-high road” model of the emotional circuit underlying associative learning^37,38^.

Since the amygdala is considered to be an evolutionary conserved structure^6^, we hypothesized that its connectivity also shows similarities across species. Thus, we next compared the network organization found in mice to human anatomical data. We applied high-resolution tractography on a large human dataset^39^ and systematically mapped connections between human homologues of LT, dMT, TeCx, and PFC and the amygdala. In line with our observations in mice, our results in humans also revealed two parallel thalamo-cortico-amygdala streams originating in the LT and MT. At the level of the neocortex, the lateral stream included the temporopolar cortex (Broadman area, BA38), while the medial stream was composed of various frontal cortical regions: the frontopolar (BA10), the orbitofrontal (BA11), the cingulate (BA24), the subgenual cingulate (BA25) and the dorsal anterior cingulate (BA32) cortices (Fig. 1f and Extended Data Fig. 4, Extended Data Video). We also revealed that MT-amygdala tracts originated in the most dorsomedial region of the MT (Fig. 1g), also called PVT in the human brain, where the presence of Calr-expressing neurons was reported earlier^23,40,41^. On the other hand, LT-amygdala tracts were most prominent in the medial pulvinar (mPul) and the medial geniculate nucleus (MGN) (Fig. 1g). Quantification of tract counts in each stream revealed that the relative strength of the lateral stream was higher at the level of thalamo-amygdala and cortico-amygdala connections, but not at the thalamo-cortical level (Fig. 1h). Interestingly, thalamo-amygdala and cortico-amygdala tracts seemingly terminated at different subregions of the amygdala. Specifically, while partially overlapping LT- and TeCx-Amy pathways terminated around the lateral (La) nuclei, most MT- and PFC-Amy pathways did so at the level of basal (Ba) and accessory basal (AB) (Fig. 1i).

Collectively, our data demonstrated a parallel architecture of two evolutionarily conserved thalamo-cortico-amygdala streams in both mice and humans. Specifically, the lateral stream originates in LT, while the medial stream arises from the dMT. These thalamic regions were connected to the amygdala both directly (∼”low road”) and indirectly (∼”high road”) by the corresponding TeCx and the PFC regions.

### Dual thalamic pathways drive different neuronal populations within the amygdala

Our macroscale data from both mice and humans suggested that the lateral and medial thalamic streams innervate the amygdala in a mostly non-overlapping manner at the level of distinct amygdala subnuclei. For a more detailed characterization of the amygdala circuitry in a reproducible and biologically relevant manner, we developed a novel immunohistochemically-based segmentation (IHCS) technique in mice to outline amygdala subnuclei (Extended Data Fig. 5-6). IHCS, using a combination of 8 different molecular markers, provided us with a precise reference framework for the localization of subsequent virus and tracer injection sites, electrode placements and axon arborizations.

To map the distribution of amygdala-projecting thalamic neurons, we performed microinjections of CTB in all major amygdala subnuclei we defined with IHCS. Only confirmed, subnucleus-selective trials were included in the analysis (Fig. 2a,b and Extended Data Fig. 7a,b). Our data (*n* = 30 mice; *N* = 14934 cells in total) revealed that amygdala subnuclei receive varying levels of thalamic innervation (Fig. 2c). By clustering the labelled neurons into 4 major thalamic territories, we revealed that the majority (65-93%) of them were located either in LT or dMT. Only a minor fraction of cells was found within the GusT or the vMT (Fig. 2d and Extended Data Fig. 7c).

**Figure 2:**
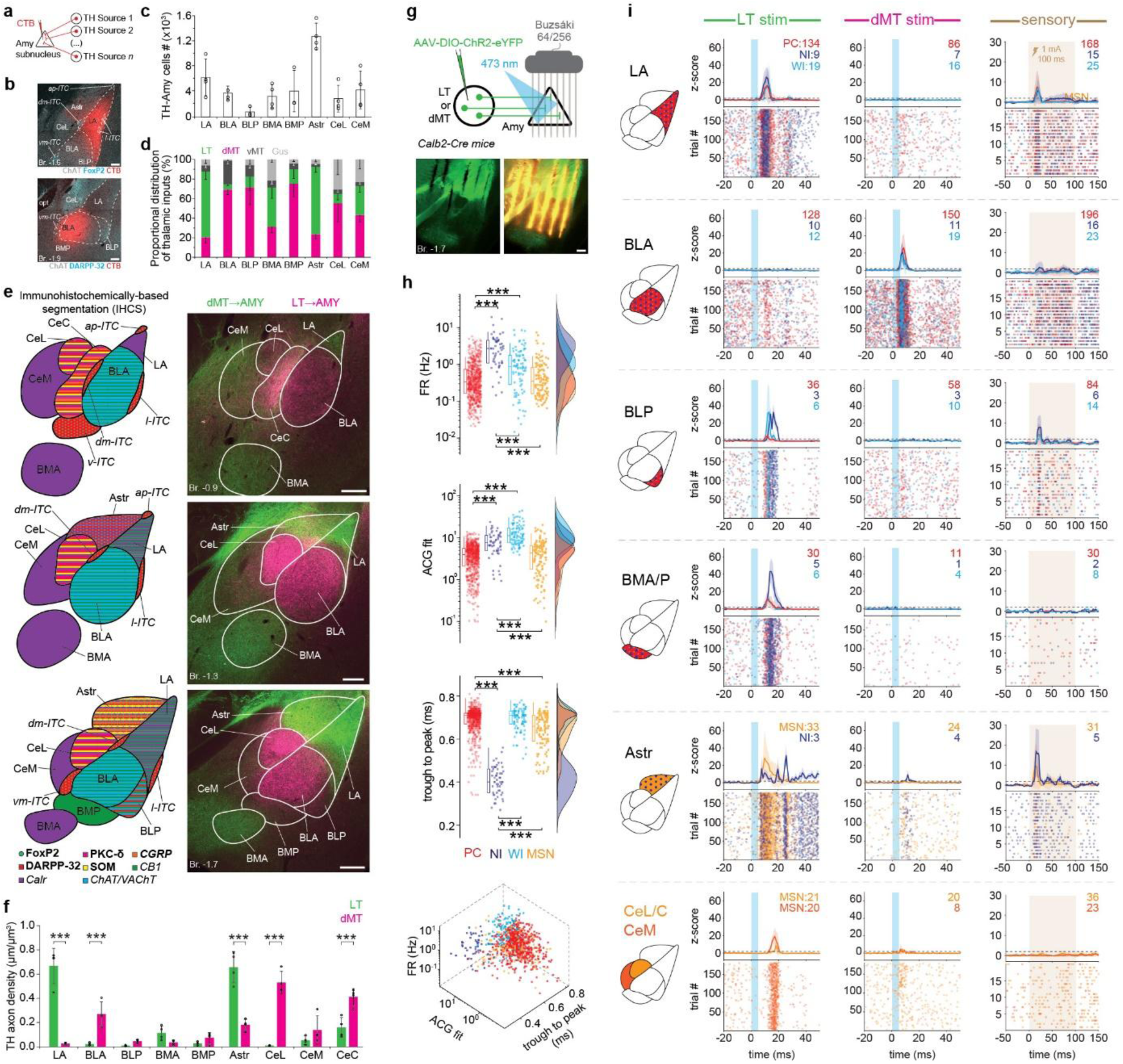
Dual thalamic pathways drive distinct neuronal populations within the amygdala. **a,** Experimental design for amygdala subnucleus-selective retrograde (CTB) tracings in the thalamo-amygdala network in mice. **b,** Representative injection sites (*top*, LA; *bottom*, BLA) confirmed using immunohistochemically-based segmentation (IHCS). For more details on amygdala subnuclei segmentation, see Extended Data Figures 5-6. Scale bars, 200 µm. **c**, Graphs show amygdala subnucleus-specific distribution of the CTB-labeled thalamic projection cells (*n* = 3-4 mice per subnucleus; one-way ANOVA, Thalamus, *F_7,22_* = 9.303, *P* = 2.404*10^-5^). **d**, Proportional graphs for each amygdala subnucleus indicate LT and dMT as the two major sources of the thalamo-amygdala innervation. **e**, Distribution of LT (magenta) and dMT (green) afferents in different amygdala subnuclei (right) identified by IHCM (left). Scale bars, 200 µm. **f**, Quantification of relative LT (green) and dMT (magenta) axon densities at the individual amygdala subnucleus level (*n* = 4 mice; one-way ANOVA, Thalamus, *F_1,8_* = 3.252*10^-4^, *P* = 0.986, Thalamus*Amygdala subnucleus, *F_1,8_* = 51.891, *P* = 1.273*10^-22^, Tukey’s post-hoc test, \*\*\**P* < 0.001). **g,** Experimental design (*top*) and histological validation (*bottom*) for high-density electrophysiological experiments. Scale bars, 200 µm. **h**, Classification of putative cell types based on firing rate (FR), autocorrelogram shape (ACG Fit) and trough to peak (*n_LT_* = 5 mice, *n_dMT_* = 3 mice; one-way ANOVA, *F(FR)_3,903_* = 89.92, *P(FR)* = 6.41*10^-51^; *F(ACG fit)_3,903_* = 165.2, *P(ACG fit)* = 2.22*10^-85^; *F(trough to peak)_3,903_* = 324.3, *P(trough to peak)* = 7.58*10^-143^; Tukey’s post-hoc test, \*\*\**P* < 0.001). 3D-plot showing clustered units separated into 4 different neuronal subtypes, pyramidal cells (PC, *n* = 633, red), narrow interneurons (NI, *n* = 56, dark blue), wide interneurons (WI, *n* = 92, light blue), and medium spiny neurons (MSN, *n* = 126, yellow). **I**, Z-score (*top*) and raster plots (*bottom*) show the spiking activity of putative neuron subtypes in distinct amygdala subnuclei evoked by LT (*left*), dMT (*middle*) and sensory stimulation (*right*). Data in (**c**,**d**,**f** and **i**) are mean ± SD. Circles in **c**,**f**,**h** and **i** indicate individual data points. For detailed statistical data, abbreviations and raw data see *Supplementary Tables* and *Source Data Tables*.

Next, we quantified the density of simultaneously labelled LT and dMT axons within the individual amygdala subnuclei by using the Calb2-Cre line (Fig. 1d). Confocal microscopic analysis revealed that overall densities of LT and dMT axons did not differ significantly at the level of the amygdala as a whole, but we found significant differences at the level of individual amygdala subnuclei (Fig. 2e,f and Extended Data Fig. 8-9). Notably, significantly more LT axons were found in the lateral nucleus (LA) and the amygdalo-striatal transition zone (Astr), an extended-amygdala region known to be involved in threat-learning related processes^16,42–44^. In contrast, the densest dMT axon arborization was found in the anterior basolateral (BLA), centrolateral (CeL) and centrocentral nucleus (CeC). Additionally, we also performed Cre-dependent AAV tracings from TeCx and PFC in Thy1-Cre mice and revealed that cortical innervation was also mostly non-overlapping, most evidently in the LA (by the TeCX) and BLA (by the PFC), indicating that cortical innervation patterns resemble their respective thalamic counterparts (Extended Data Fig. 10-12). Collectively, our anatomical experiments revealed that the LT and dMT provide the vast majority of the thalamo-amygdala innervation, and – together with their corresponding cortical regions – they selectively target different amygdala subnuclei in a dominantly non-overlapping manner.

To characterize the functionality of these thalamo-amygdala streams, we injected Cre-dependent AAV expressing ChR2, unilaterally into the LT or dMT of Calb2-Cre mice and performed in vivo acute electrophysiological recordings. We used high-density silicon probes covering the whole amygdala to capture neuronal responses evoked by optogenetic or sensory stimulation within different amygdala subnuclei simultaneously (Fig. 2g). We utilized the recently developed CellExplorer tool^45^ to create a large dataset of identified single unit clusters (907 in total). Then, these clusters were separated into different neuronal subtypes based on their location, firing rate (FR), autocorrelogram shape (ACG Fit) and trough to peak parameters: pyramidal cells (PC), interneurons with narrow spikes (NI), interneurons with wide spikes (WI) and medium spiny neurons (MSN; Fig. 2h). Here, we also used our previously established IHCS framework to assign our clusters to subnucleus-specific subgroups, by reconstructing the location of recording sites registering the highest amplitude action potential for each individual putative neuron. Note that CeL and CeC, as well as BMA and BMP recording sites could not be separated reliably (see Methods), so we treated them together. In general, optogenetic stimulation of LT and dMT axons in the amygdala elicited markedly different responses across individual amygdala subnuclei (Extended Data Fig. 13a and Supplementary tables). In line with our anatomical results, we confirmed that LT axonal stimulation induced robust neuronal activation with short latency (<10 ms) in LA and Astr, while dMT axons only triggered a similarly robust and short latency response in the BLA. We also recorded significant neuronal activation during LT stimulation from BLP, BMA/P, CeL/C and CeM subnuclei with a longer latency (>10 ms), while comparable responses were absent during dMT stimulation. Interestingly, Astr activity had multiple peaks in the first 30ms implicating the existence of direct and indirect thalamic drives (Fig. 2i and Extended Data Fig. 13b). We also applied footshocks to investigate activity patterns evoked by an exogenous, affective sensory stimulus. In this case, we found that robust and short latency activations (<20 ms) were also present in LA and Astr, while responses with longer latency (>20 ms) were evoked in BLP (Fig. 2i and Extended Data Fig. 13a,b). These results demonstrated that direct optogenetic stimulation of the LT pathway and sensory stimulation evoke similar neuronal activity patterns within the amygdala, while the excitation of dMT inputs only evoked direct responses in the BLA region, without any subsequent intra-amygdala propagation. Collectively, our anatomical and physiological data highlighted that the lateral and medial thalamo-amygdala streams drive distinct neuronal populations and their selective activation results in contrasting activity patterns within the amygdala.

### LT and sensory inputs drive divergent intra-amygdala activation

Our electrophysiological data with optogenetic and sensory stimulations showed evoked activity patterns with distinct spatial and temporal distribution within the amygdala. To further investigate the temporal processing dynamics of intra-amygdala activity at the subnucleus and single unit level, we calculated the cumulative sum of spikes within a 30 ms pre- and post-stimuli time window, which allowed us to identify time-points with significantly increased neuronal activation (Fig. 3a, cumulative mean above a threshold of significance). We found that LT-inputs elicited fast LA and Astr response (8 ms) and the LA activation likely propagated through monosynaptic connections (within 1-5 ms)^46,47^ towards the BLP/BMP (12 ms) and subsequently, towards the CeL/C and CeM (15-19 ms) subnuclei. In more details, LT stimulation elicited a distinct multi-stage intra-amygdalar activation pattern, initiated by rapid-onset responses in the LA and Astr. These initial signatures were consistent with feed-forward inhibition^16^, evidenced by the highly synchronous recruitment of putative PCs, NIs, MSNs, and WIs. This primary wave was followed by the downstream recruitment of the BLP and BMA/P, which exhibited characteristic inhibitory feedback motifs with distinctively lagging NI peaks. Finally, late-phase activation of CeL/C and CeM putative MSNs, alongside a secondary wave of Astr NI activity, likely represents the terminal gating mechanism regulating the functional output of the network. In contrast, dMT-inputs only evoked fast responses within the BLA (6 ms), with a minor downstream effect in the Astr. Notably, sensory stimulation elicited an activity pattern qualitatively similar to the optogenetic stimulation of LT-inputs in the LA, BLP and Astr, which was also confirmed by comparing the response latency of individual neurons. The activation probability and proportional latency distributions of unit responses also demonstrated a comparable, sequential activity pattern within the amygdala (Fig. 3b and Extended Data Fig. 13b). To further validate these findings, we correlated normalized thalamic axon density values from the experiments discussed earlier (Fig. 2e,f) with the magnitude of thalamic evoked early (putative monosynaptic) responses at the subnucleus level within the amygdala. Confirming our results, we found that thalamic stimulation elicited stronger responses in those amygdala subnuclei that received anatomically more robust innervation from the same thalamic source (Fig. 3c,d). In case of the dMT-linked amygdala circuit, significant correlation was only found without including data from the CeL/C due to the high axon density but low evoked response values (Fig. 3c and Extended Data Fig. 14). This discrepancy can be explained by the thalamic neurotransmitter (the brain-derived neurotrophic factor) released in the CeL having slow kinetics^13^ (also see the gradually built-up responses after ∼ 100 trials).

**Figure 3:**
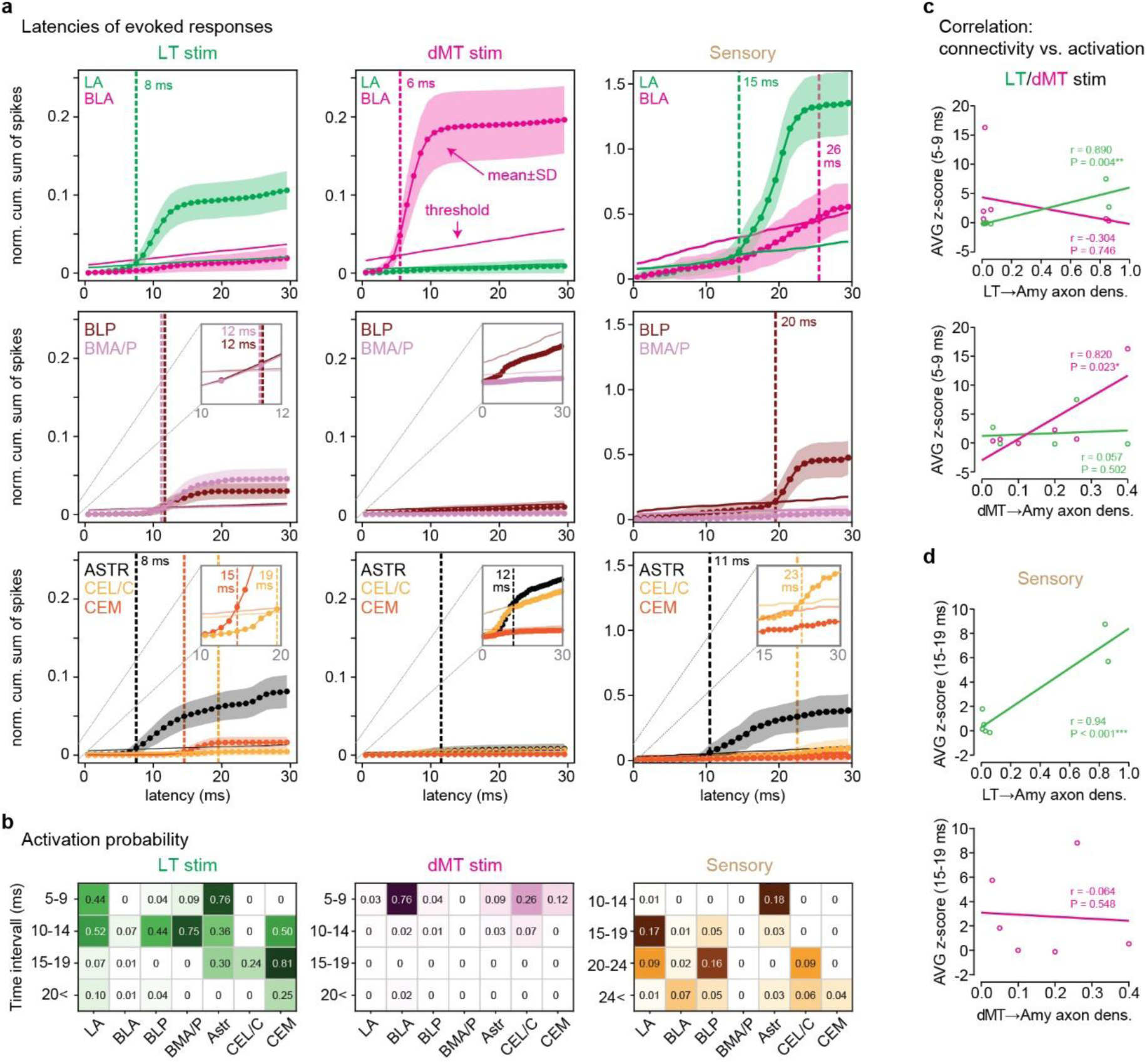
LT and sensory inputs evoke sequential amygdala responses in contrast to a discrete activation evoked by dMT. **a**, Spiking activity pooled from all neuronal subtypes within a subnuclei (the same dataset as in Fig. 2i). Dotted solid lines with shaded area indicate normalized mean ± SD cumulative sum of spikes (0 to 30 ms, post-stimulus). Single solid lines indicate the threshold of significance for activation (mean normalized cumulative sum + SD of baseline activity, calculated from -30 to 0 ms, pre-stimulus). Vertical dashed lines indicate the latency of significant activation in given subnuclei (positive threshold crossings). **b**, Activation probability of neurons within different amygdala subnuclei in response to LT, dMT or sensory stimuli. **c**,**d**, Correlations between the thalamo-amygdala anatomical (normalized axon density; x axis) and functional connectivity obtained with optogenetic (**c**) and sensory (**d**) stimulation [(analyzed as the averaged z-score calculated in a given post-stimulus 5-9 (optogenetic) and 15-19 ms (sensory) interval; y axis; Pearson correlation analysis, \**P* < 0.05, \*\**P* < 0.01)]. Note the dMT panels are without CeL/C (for data including CeL/C, see Extended Data Fig. 14). Circles indicate anatomical and electrophysiological value pairs for each amygdala subnucleus. For detailed statistical data see *Supplementary Tables*.

Taken together, LT stimulation gave rise to a multi-step intra-amygdala activity pattern initiated by a direct, short-latency LA activation, followed by the activation of downstream cortical (BLP, BMA/P) and finally, striatal (Astr, CeL/C, CeM) subnuclei. In contrast, dMT stimulation elicited a strong short-latency response only in the BLA, without any further propagation to other amygdala subnuclei.

### LT-activated LA forms parallel intra-amygdala pathways

In order to anatomically confirm the intra-amygdala connectivity patterns suggested by our electrophysiological data, we applied complementary tract-tracing methods in a subnucleus-specific manner. First, after placing micro-deposits of CTB into each amygdala subnucleus (Fig. 2a,b), we quantified the number of retrogradely labelled cells in the other 7 amygdala subnuclei (15626 cells in total). In general, our data uncovered that individual subnuclei received a distinct combination of inputs with varying strength from other amygdala subnuclei (Fig. 4a). In accordance with our electrophysiological findings, the most prominent downstream targets of the LA were the Astr, BLP and BMP. BLP and BMP, in turn, strongly innervated the central nuclei (CeL/C, CeM), as well as the Astr (Fig. 4a,b and Extended Data Fig. 15). In addition, CeM was also strongly connected by the BMA. On the other hand, we found that BLA was not a major source of innervation in any other amygdala subnucleus (Fig. 4b).

**Figure 4:**
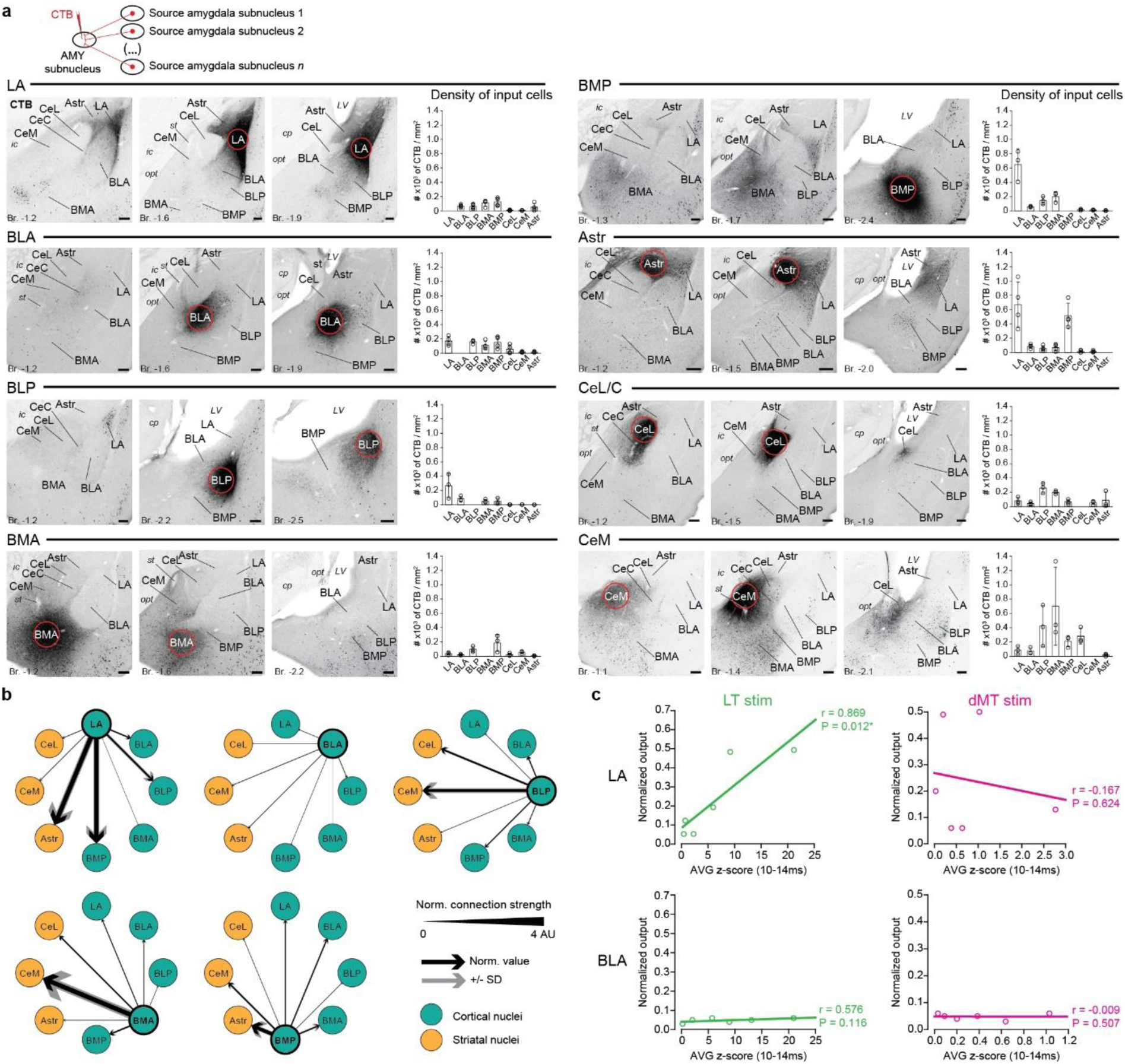
LT-activated LA forms parallel intra-amygdala pathways. **a**, Distribution of CTB-labeled input cells within amygdala subnuclei to LA, BLA, BLP, BMA, BMP, Astr, CeL/C and CeM. *Top*: experimental design for subnucleus-specific retrograde tracing. *Left*, 3 representative images from each trial with the CTB injection sites (red circles) and the labeled CTB neurons (small black dots). *Right*, quantification for input strengths as densities of CTB-labeled cells. Data in are mean ± SD and circles indicate individual data points. Scale bars, 200 µm. **b**, Vector diagrams showing efferent connectivity strengths for each subnucleus (normalized data from **a**). Thickness of the arrows is scaled from 0 to 4 normalized values. Black lines represent mean, gray outline indicate SD. **c,** Correlations between normalized intra-amygdala anatomical (CTB; y axis) and functional connectivity (analyzed as the averaged z-score calculated in the 10 to 14 ms post-stimulus interval, x axis; Pearson correlation analysis, \**P* < 0.05). Circles indicate anatomical and electrophysiological value pairs for each amygdala subnucleus. For detailed statistical data, abbreviations and raw data see *Supplementary Tables* and *Source Data Tables*.

We also used correlational analyses to validate our intra-amygdala anatomical and electrophysiological findings. In order to detect disynaptic propagation, we quantified the magnitude of optogenetically evoked responses (z-score) within the post-stimulus interval (10-14 ms) averaged for each amygdala subnucleus. This data was then correlated with normalized intra-amygdala CTB-labelled cell densities, revealing that the LT-induced late activation pattern showed strong correlation with the intra-amygdala connectivity pattern of LA (Fig. 4c) which subnucleus itself was directly and strongly innervated by the LT (Fig. 2). Since BLA was found to be a negligible source of intra-amygdala innervation and dMT stimulation elicited only short-latency activation in the amygdala, we could not detect significant correlation between the anatomical connections of BLA and dMT-induced late activity patterns (Fig. 4c). Collectively, our anatomical and electrophysiological observations uncovered that the LT-driven LA is the main source of intra-amygdala information flow. In contrast, the dMT-innervated BLA only weakly participates in the signal processing within the amygdala.

### Anterograde tracing evidence for the LA-linked intra-amygdala connectivity patterns

Besides the retrograde tracing we also conducted subnucleus-selective anterograde tract-tracing approaches to further evaluate the intra-amygdala wiring. We performed iontophoretic microinjections of anterograde tracer biotinylated dextran amine (BDA) into the cortical-like subnuclei and found a connectivity pattern matching our retrograde tracing data. Specifically, LA innervates BLP, BMP and Astr and, in turn, BLP is strongly connected to the central subnuclei, while BMP to the Astr. On the other hand, BLA provides only a sparse innervation to any other amygdala subnucleus (Fig. 5a,b). These data confirm that LA but not BLA neurons participate in the intra-amygdala wiring. Furthermore, besides the direct LA-Astr route, the LA-connected BLP and BMP neurons provide the major striatal (CeL/C, CeM and Astr) outputs of the cortical amygdala.

**Figure 5:**
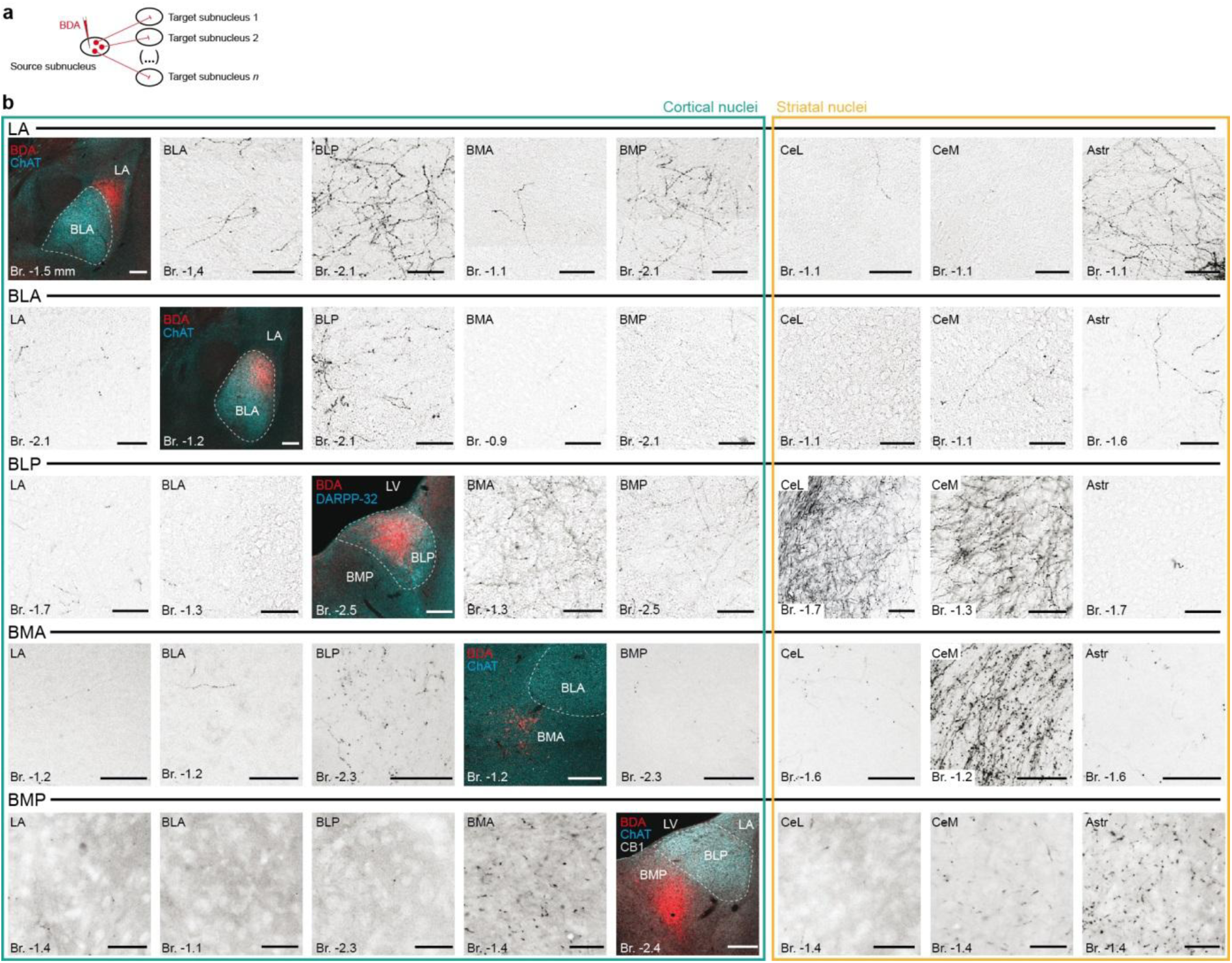
Anterograde tracing data confirm the unexpected connectivity motifs within the amygdala complex. **a**, Experimental design for subnucleus-selective anterograde tracings (biotinylated dextran amine; BDA). **b**, Representative trials for output (efferent) connectivity of each cortical amygdala subnuclei. The selectivity of each injection site has been confirmed by IHCS (*fluorescent panels*). Representative high magnification images show the axonal projection densities for each trial in other amygdala subnuclei. Scale bars, brightfield images: 50 µm, fluorescent images: 200 µm.

### LA has strong intra-amygdala, while BLA has strong extra-amygdala connectivity

Our findings so far implied that the two major cortical-like amygdala subnuclei, the LA and BLA possess markedly distinct afferent and efferent connections. To further confirm this notion with direct comparison, we turned to an intersectional approach to selectively label larger number of LA and BLA neurons. As LA neurons preferentially innervate TeCx, while the BLA neurons target PFC (Extended Data Fig. 16), we injected retrogradely transporting Canine adenovirus type 2 carrying the Cre-recombinase gene (CAV2-Cre) into the TeCx or PFC of wild type mice, and Cre-dependent AAV-DIO-mCherry into the amygdala to specifically label LA or BLA neurons, respectively (Fig. 6a,b). Connectivity mapping confirmed that LA neurons have strong intra-amygdala but weak brain-wide (extra-amygdala) efferentation, while the connectivity profile of the BLA showed the opposite: strong innervation to various brain regions outside the amygdala, such as the PFC, dorsal striatum (CPu) and nucleus accumbens (NAc), bed nucleus of the stria terminalis (BNST) or the claustrum (CL; Fig. 6a,c and Extended Data Fig. 17). We further confirmed these findings with classical anterograde tracing and Cre-dependent AAV tracing in the *Arhgef6*-Cre (also known as NL189-Cre) mouse line by labelling primarily BLA neurons (Extended Data Fig. 18)^48^. In sum, we confirmed our electrophysiological findings and retrograde intra-amygdala connectivity analysis with various anterograde tracing techniques and also described the extra-amygdala connectivity of LA and BLA neurons.

**Figure 6:**
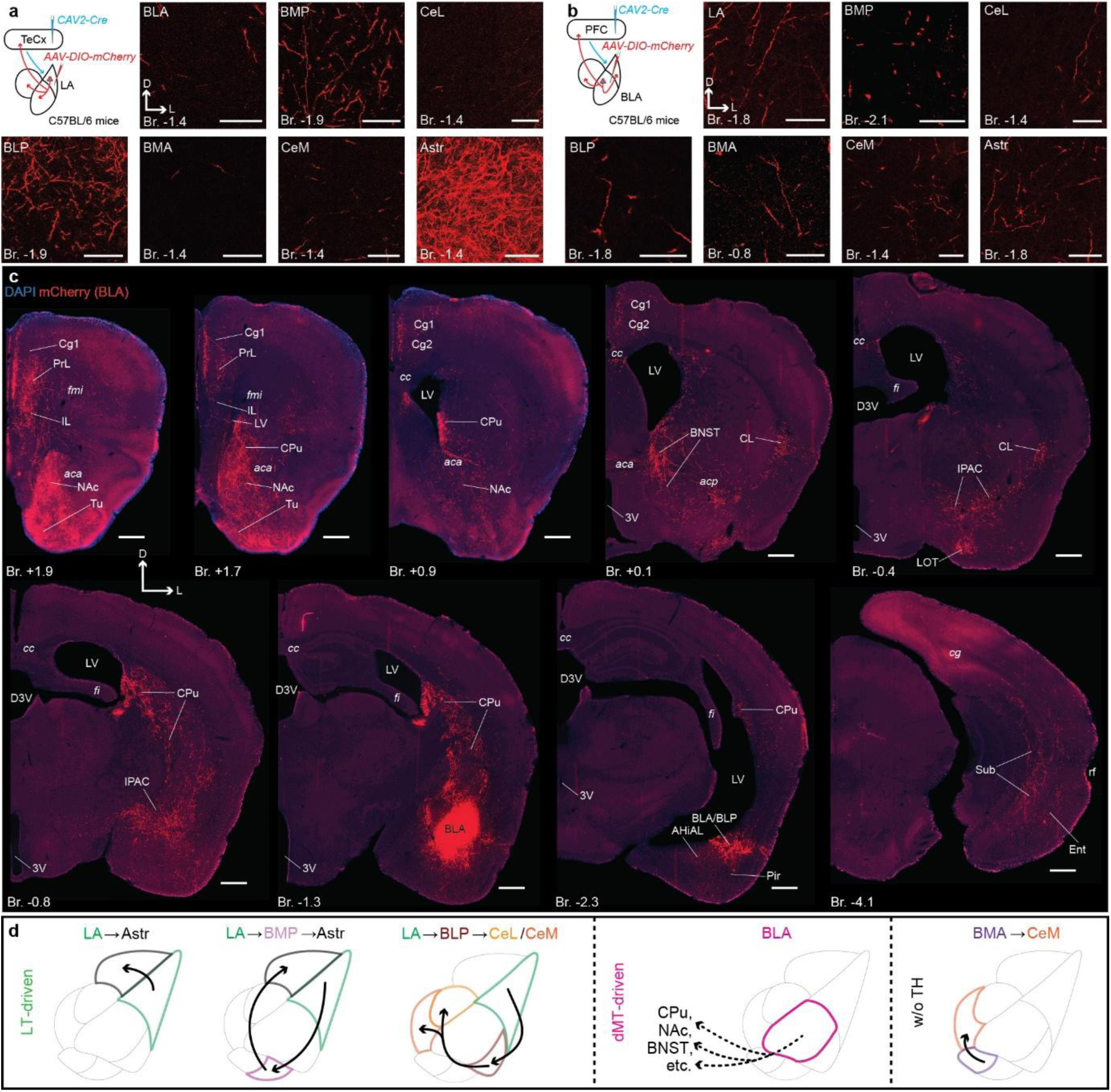
LA has strong intra-amygdala connectivity, while BLA has strong extra-amygdala connectivity. **a**, Experimental design and a representative trial for CAV2-Cre-mediated viral tracing of LA. High magnification images show the axonal projection densities in BLA, BLP, BMA, BMP, Astr, CeL, CeM. Scale bars, 50 µm. **b**, Experimental design and a representative trial for CAV2-Cre-mediated viral tracing of BLA. High magnification images show the axonal projection densities in other amygdala subnuclei. Scale bars, 50 µm. **c**, Representative overview images (of the same trial as in **b**) show strong extra-amygdalar axonal projections of BLA in various brain regions including the prefrontal cortex (Cg1, PrL, IL), dorsal (CPu) and ventral striatum (NAc), bed nucleus of the stria terminalis (BNST) and the claustrum (CL), among others. Scale bars, 500 µm. For abbreviations see *Supplementary Tables*. **d**, Summary diagram for the main pathways described here using subnucleus-selective tracings and electrophysiological recording. Note that (i) three major intra-amygdala routes originate from the LT-driven LA; (ii) the dMT-targeted BLA is strongly connected to various extra-amygdala targets, but not to the other amygdala subnuclei; (iii) BMA, receiving relatively weak or no thalamic innervation, mostly targets the CeM.

Collectively, our data demonstrate that the thalamo-cortico-amygdala circuits originate in Calr-expressing thalamic neurons and form two major streams. Specifically, LT is positioned to drive a multi-step intra-amygdala network via the LA, while dMT innervation of the BLA serves as a link towards various limbic brain regions outside the amygdala.

## Discussion

Despite being fundamental elements of the emotional processing circuitry, the way the amygdala computes the incoming signals as well as the way these drive intra-amygdala pathways, have not been completely characterized so far. In this paper, we report a previously overlooked complexity in the amygdala circuitry, mainly defined by two, mostly non-overlapping thalamo-amygdala pathways originating from Calr-expressing thalamic neurons in mice. Specifically, LT neurons predominantly innervate the LA which, in turn, drives an interconnected intra-amygdala network involving several other amygdala nuclei, ultimately terminating in striatal regions such as the Astr, CeL/C and CeM. We also showed that this circuitry is involved in the processing of peripheral sensory information (pain), in accordance with previous reports^16,49^. On the other hand, dMT axons are most abundant in the BLA where they drive a prominent short-latency response. However, the BLA outputs are directed towards extra-, rather than intra-amygdala targets, such as the PFC, NAc, BNST or the CL, providing a link to a brain-wide saliency network. This non-overlapping duality was also noticeable at the level of cortico-amygdala connections^34–36^, as well as in our human tractography data.

As a general remark, it is important to stress that the lack of a commonly accepted and experimentally reproducible delineation approach to separate amygdala nuclei can obstruct anatomically precise investigations. Additionally, inconsistent nomenclatures, and abbreviations in particular, present in the most widely used anatomical atlases, review papers and original research articles can be the source of further confusion and misunderstandings in the field (see Supplementary Discussion for details). The IHCS approach we proposed to delineate amygdala subnuclei in mice (Fig. 2. and Extended Data Fig. 5-6) could be a universal tool to overcome such inconsistencies to produce reproducible results. This approach provided us with the opportunity to gather data with fundamentally different experimental techniques like anatomical tracing and *in vivo* electrophysiology, while keeping a high standard of anatomical precision, comparability and reproducibility.

Our results demonstrated that Calr-expressing neurons in the LT and dMT provide the majority of input in the amygdala in a non-overlapping manner, clarifying prior findings^21,22,49–51^. In fact, our earlier studies independently reported that Calr is a specific molecular marker for LT^16^ and dMT^23^ neurons. In recent years certain functions were linked to both the LT- and dMT-amygdala pathways. For instance, both the LT and the dMT were implicated in plastic fear memory formation via the amygdala^12,16,18,52^. Similarly, arousal regulation was also identified as a function of both the LT and the dMT^23,53–56^. However, the clear anatomical separation of these pathways reported here, as well as the fact that peripheral sensory stimulation failed to elicit significant electrophysiological responses in the BLA indicate that some aspects of emotionally relevant information are processed differently in these systems.

We also described a comparable anatomical duality in the human thalamo-cortico-amygdala network, confirming previous studies^39,57,58^. Our results also indicate that the relative weight of the LT and TeCx pathways to the amygdala is higher in the human brain. An evolutionary increase in the functional importance of these pathways would not be surprising, considering the prominent expansion of the human pulvinar and TeCx (Bartlett, 2013; Homman-Ludiye and Bourne, 2019; Braunsdorf et al., 2021). Connections between the PFC and pulvinar, as well as between the TeCx and the medial thalamus, have been reported in previous primate anatomical^61,62^ and human functional connectivity studies^63,64^. Interestingly, we found minimal cross-connectivity between the lateral and medial streams at the level of cortico-thalamic connections in the mouse brain. Collectively, these results suggest that although some fundamental wiring patterns in the amygdala network remained evolutionarily conserved in the human brain, notable differences might have emerged as well.

With the combination of anatomical tracing and high-density electrophysiological recordings, we revealed an overlooked connectivity pattern of individual amygdala nuclei. Intriguingly, although connections within the amygdala have been extensively investigated in the previous decades, even recent review articles present different variations of intra-amygdala wiring maps^10,28,65^. While the exact anatomical weight and functional significance of LA-/BLA-to-CeL connections in the fear circuitry have been questioned by some authors^27,66^, these connections consistently receive disproportionally more attention than other, anatomically better documented amygdalo-striatal streams, such as the LA-to-Astr^16,67–70^ or BLA-to-NAc pathways^29–31,71–74^. We also found strong LA-Astr and BLA-NAc connections, but only weak-to-absent innervation to the CeL/C from either nucleus, in accord with previous anatomical^67,68,75–77^, electrophysiological^69,70,78^ and functional reports^79^. We only found significant innervation to the CeL/C from the posterior-lateral aspects of the basolateral amygdala – that we refer to as BLP -, resembling earlier findings^31–33^. On the other hand, we showed direct anatomical and functional connection between the LA and BLP, indicating that information from the LA can ultimately reach the CeL/C and CeM indirectly, perhaps integrating additional information from other sources, such as the rhinal cortices or the HPC via the BLP along the way^34,80^. Furthermore, we also demonstrated a prominent indirect pathway from the LA to the Astr and CeM through the BMP, which can be an additional point of integration for prefrontal innervation^34,81^. In sum, LT-driven LA neurons are positioned to drive multiple parallel intra-amygdala pathways towards the Astr, CeL/C and CeM, and, at the same time, providing entry points for incoming cortical information from various sources that otherwise do not reach the LT or LA directly.

Our results also suggest that the BLA plays the role of connecting the dMT to a broad network of limbic regions, rather than being a key element of the intra-amygdala network. In accord, extra-amygdala connections of the BLA, such as the PFC, NAc or HPC^34^, were implicated in various positive and negative emotional behaviors^29–31,82,83^. Interestingly, the dMT itself also innervates many BLA targets directly, such as the PFC, NAc or BNST^22,23^, suggesting that the BLA might add an extra layer of information processing to the circuitry linked to the dMT. In fact, BLA receives innervation from the dorsal raphe^84^, CL^85^ and ventral tegmental area^86^, among others, that do not reach the dMT directly. Since both the dMT and the BLA were implicated in the regulation of arousal, saliency detection and action selection^14,23,87–90^, it can be postulated that the dMT-BLA pathway is part of a brain-wide emotional state-setting network, contributing to fear-related behavior among other functions^91,92^. Collectively, our results depict the amygdala as a complex network of interconnected, but also parallel pathways involved in various operations, rather than single processing unit with a simple, linear information flow.

In sum, we demonstrated that LT is positioned to drive a complex network of parallel intra-amygdala pathways originating in the LA and terminating in known striatal output regions of the amygdala, namely the Astr, CeL/C and CeM. We also demonstrated that this thalamo-amygdala pathway can be driven by naturalistic peripheral sensory stimulation. These pathways can also link information originating in the LT with cortical input from various sources. On the other hand, dMT innervation in the BLA seems to serve as a link towards a more widespread network, ideally positioned to contribute to state-setting during emotional behavior. Further behavioral experiments could reveal the functional consequences of this duality in the amygdala circuitry, possibly helping us understand the mechanisms underlying various emotional regulations.

## Supplementary discussion

Although the amygdala is one of the most intensively investigated regions in the brain, a single, universally accepted approach to its parcellation has not been implemented yet. Focusing on the mouse brain, even the most popular anatomical atlases, such as The Mouse Brain in Stereotaxic coordinates^93^ or the Allen Mouse Brain Reference Atlas^94^ (https://atlas.brain-map.org/) present notable differences (see also Extended Data Fig. 5). Further complicating the interpretation and comparison of results, abbreviations are even less consistent across rodent studies and atlases, even in relatively recent publications. For example, some studies label the (anterior and posterior) basolateral nuclei with the abbreviation “BA”^95,96^ or “BL”^97^, while others use the “BA” abbreviation for the “basal amygdala” collectively referring to the anterior and posterior basolateral and basomedial nuclei^98–100^. In contrast, other authors refer to the lateral and basolateral nuclei collectively as “BLA”^30,31,81,101–103^ or “BLC”^104^, while others include all the above-mentioned cortical-like amygdala nuclei in the basolateral complex also abbreviated as “BLA”^25,105,106^. Further, notably different abbreviations can also be found in papers published earlier^79,107–109^. Additionally, simply using terms such as “basolateral amygdala” or “basolateral complex of the amygdala” without a specific definition is also not uncommon.

Anatomical precision is particularly important in the amygdala, due to its relatively small size and ventral position in the brain which makes its investigation physically challenging, and also due to its functionally different, yet anatomically intermingled subdivisions. For instance, in accordance with other reports^110–114^, we demonstrated that the neighbouring Astr and CeL have fundamentally different intra- and extra-amygdala connections. Considering that the CeL is usually approached from the dorsal direction (i.e. through the Astr) with injection pipettes, electrodes or optical fibers, precise differentiation of CeL and Astr would be crucial. Similarly, the LA and BLA are also situated in a neighbouring manner, without any apparent physical barriers (e.g., white matter bundles) between the two nuclei. Nevertheless, many publications fail to report the way the authors separated LA and BLA investigations. Additionally, injection sites, probe or optical fiber placements are often not shown in publications or shown in a way that makes the identification of the exact location challenging (e.g., without background staining or stereotaxic coordinates). Although a few genetic markers have been used to label different subsets of neurons in these nuclei, none of these became generally implemented in the field^32,48,115^.

Such inconsistencies and the lack of commonly accepted anatomical standards can seriously obstruct advances in the field of amygdala research, especially in the age of search engines and large language models when the significance of abbreviations, keywords and search phrases, as well as the rate of information proliferation increases. In this publication, we consistently used a nomenclature and abbreviations mostly matching the 5^th^ edition of The Mouse Brain in Stereotaxic Coordinates^93^, based on our IHCS amygdala parcellation (Extended Data Fig. 5-6).

## Methods

### Animals

Adult (3-5 months old, male and female) C57BL/6j, *Thy1*-Cre (FVB/N-Tg(Thy1-cre)1Vln/J, RRID: IMSR_JAX:006143), *Calb2*-Cre (B6(Cg)-Calb2^tm1(cre)Zjh^/J, RRID:IMSR_JAX:010774; The Jackson Laboratory, Bar Harbor, Maine, USA) and NL189-Cre (Tg(*Arhgef6*-cre)NL189Gsat/Mmucd) were used for the experiments. They were group housed in a humidity- and temperature-controlled environment. Animals were entrained to a 12 h light/dark cycle (light phase from 07:00 AM) with food and water available *ad libitum*. All experiments involving experimental animals were carried out in the Research Centre for Natural Sciences and the Institute of Experimental Medicine of the Hungarian Research Network and were officially approved by the Regional and Institutional Committees of both Institutes. The experiments were also approved by the National Animal Research Authorities of Hungary (PE/EA/00080-4/2023).

### Stereotactic surgeries

Animals were anesthetized under ketamine-xylazine (100 mg/kg; 4 mg/kg; i.p.) during surgeries involving tracer or viral injections. After losing consciousness, animals were fixed to the stereotactic apparatus (RWD Life Sciences, Kopf Instruments) with a mouse-specific ear-bar. Lidocaine (100 mg/ml) was sprayed in the ear canal for analgesia. Respiration was monitored during surgeries and a heating pad was set to 37°C to avoid hypothermia. After removing the hair from the scalp, the skin was cleaned with saline and Betadine and the surgical area was sprayed with Lidocaine (100 mg/ml). The skin was then incised and the skull surface was cleaned with saline. Craniotomies at the appropriate coordinates (see below) were made using a dental drill. After tracer/virus injections, the skin was sutured and the wound was treated with Betadine ointment. After surgeries, animals received Rimadyl injections (Carprofen, 1.4 mg/kg; i.p.) for analgesia and additional saline (i.p.) to avoid dehydration. Animals were allowed to regain consciousness on the heating pad before returning them to their homecage. Paracetamol (0.3 mg/ml) in drinking water was provided *ad libitum* for 2 days after the surgery for additional analgesia.

### Classical tracing experiments

Retrograde tracing was carried out with 0.5% Cholera Toxin B subunit (CTB; List Biological Laboratories: 104) injected into different amygdala subnuclei (for coordinates see Supplementary Tables). For anterograde tracing 5% biotinylated dextran amine was used (BDA, MW: 10.000, Molecular Probes: D1956, RRID: AB_2307337). Tracers were iontophoretically injected (CTB: 7-7 sec on/off duty cycle, 3-5 µA, 5-10 min; BDA: 3-3 sec on/off duty cycle, 2 µA, 3-5 min) with IonFlow Bipolar electrophoretic equipment (Supertech Instruments, Hungary). Animals were perfused after a 7-10 days of survival time under ketamine-xylazine-pipolphen (10 mg/kg; 50 mg/kg; 5mg/kg; i.p.) anesthesia, first with cold (4°C) 0.9% NaCl (Sigma-Aldrich, CAS No.: 7647-14-5) solution, then, with ∼150 ml of cold 4% PFA (Sigma-Aldrich, CAS No.: 30525-89-4) in 0.1 M phosphate buffer (PB). Post-fixation in 4% PFA solution was done for 1-2 hours at room temperature.

### Viral injections

For cell type-specific anterograde viral tracing, AAV5-EF1a-DIO-eYFP-WPRE-hGH (30-100 nl; Penn Vector Core; #27056-AAV5; titer: 5*10^12^ GC/ml; AAV-DIO-eYFP in figures), AAV5-EF1a-DIO-mCherry (30-100 nl; UNC Vector Core; #50462; titer: 7*10^12^ GC/ml; AAV-DIO-mCherry in figures), AAV5-EF1a-dflox-hChR2(H134R)-mCherry-WPRE-hGH (50-100 nl; Penn Vector Core; #20297-AAV5; titer: 7.35*10^12^ GC/ml; Cre-ON in figures) or AAV2/8-hSyn1-DFO-hChR2(H134R)-eYFP (50-100 nl; Cre-OFF in figures)^16^ were injected at a rate of 0.5-1 nl/sec into dMT, LT (Calb2-Cre), PFC, TeCx (Thy1-Cre) or BLA (NL189-Cre) (for coordinates see Supplementary Tables) using Nanoliter Injector (World Precision Instruments, FL, USA). For electrophysiological recordings, AAV5-EF1a-DIO-hChR2(H134R)-eYFP-WPRE-hGH (50 nl / depth; Addgene 20298) was injected at a rate of 2 nl/sec into dMT or LT (Calb-Cre). Animals used for anterograde tracing experiments were perfused after 4-6 weeks of survival time. Viral expressions were always analyzed after immunohistochemical-enhancement^116^, including eYFP.

### CAV2-Cre mediated viral tracing

In order to selectively label LA and BLA principal neurons, based on the connections with the TeCx and PFC, respectively, we injected Canine adenovirus type 2 carrying Cre-recombinase gene (CAV2-Cre, CMV promoter, 50-100 nl, titer: 2.5*10^10^ pp/ml, Plateforme de Vectorologie de Montpellier, France) into the TeCx or PFC of C57BL/6j animals. During the same surgery AAV5-EF1a-DIO-mCherry (40-50 nl; UNC Vector Core; #50462; titer: 7*10^12^ GC/ml) was injected into the LA or BLA, respectively. All injections were made using a Nanoliter Injector (World Precision Instruments, FL, USA). Animals were perfused after 4 weeks of survival time.

### Tissue processing for immunohistochemistry

For prolonged periods tissue samples were stored in 0.05% sodium-azide (Sigma-Aldrich, CAS No.: 26628-22-8) solution at 4°C or in 30% sucrose (Sigma-Aldrich, CAS No.: 57-50-1) - 30% ethylene glycol (Sigma-Aldrich, CAS No.: 107-21-1) solution at -20°C. Tissue blocks were cut on a VT1200S Vibratome (Leica) into 50 μm coronal sections. Free-floating sections were intensively washed with 0.1 M PB. All antibodies were diluted in 0.1 M PB. For immunohistochemical labeling, all sections were first treated with a blocking solution containing 10% normal donkey serum (NDS, Sigma-Aldrich: S30-M) or 10% normal goat serum (NGS, Vector: S-1000, RRID: AB_2336615) and 0.5% Triton-X (Sigma-Aldrich, CAS Number: 9036-19-5) in 0.1 M PB for 30 minutes at room temperature (RT).

### Fluorescent immunohistochemistry

After the blocking solution, sections were incubated in primary antibody solution (for details see Supplementary Tables) overnight at RT or for 2-3 days at 4°C. After primary antibody incubation, sections were intensively washed in 0.1 M PB, then incubated in secondary antibody solution (for details see Supplementary Tables) for 2h at RT.

When necessary, staining was enhanced after primary antibody incubation with biotinylated secondary antibodies (1.5h, RT), Elite Avidin-Biotin Complex (eABC, 1:300, Vector Laboratories: PK-6100, RRID: AB_2336819; 1.5h, RT) and streptavidin conjugates (2h, RT; for details see Supplementary Tables). Intensive washing in 0.1 M PB was carried out after each incubation. All fluorescent slices were mounted in Vectashield (Vector Laboratories: H-1000, RRID: AB_2336789; with DAPI: H-1200, RRID: AB_2336790) and stored at 4°C.

### Immunoperoxidase staining

For the quantification and plotting (see below) of retrogradely labelled neurons, and for the visualization of labelled axons we used immunoperoxidase staining and with nickel-amplified 3-3’-diaminobenzidine (DAB; Sigma-Aldrich; CAS Number: 91-95-2) technique (DAB-Ni). Samples were treated first with 1% H_2_O_2_ solution for 10 minutes to block endogenous peroxidase activity. Then, after intensive washing, we used 10% NDS and 0.2% Triton-X solution as a blocking serum (30 mins, RT). After primary antibody incubation (CTB, GFP, mCherry, see above), slices were incubated in biotinylated secondary antibody (bHAG, bGAR) and eABC (see above). Then we developed DAB-Ni for 4-6 minutes. Sections were then dehydrated in xylol (2*10 mins) and mounted in DePex (Serva, Heidelberg, Germany; Cat. No.: 18243).

### Microscopy

Fluorescent sections were first analyzed with epifluorescent microscope (Leica DM 2500, Leica Microsystems GmbH; Camera: Olympus DP73, CellSens Entry 1.16, Olympus Corporation) at lower magnification (2.5x N PLAN 2.5x/0.07 ∞/-/OFN25, 5x HCX FL PLAN 5x/0.12 ∞/-/B) to localize injection sites and labelled neurons. Higher magnification (10x Plan Apochromat 10x/0.45 M27; 20x Plan Apochromat 20x/0.8 M27; 63x Plan Apochromat 63x/1.4 Oil DIC M27; 10x Plan Fluor 10x/0.3; 20x Plan Apo VC 20x/0.75) images were taken with a confocal microscope (Zeiss LSM 710; Zeiss ZEN 2010B SP1 Release version 6.0; Carl Zeiss Microimaging GmbH; Nikon Ni C2; NIS Elements AR, Version: 4.51.01 (Build 1146); Nikon Europe BV). Brightfield imaging, analysis of the distribution of retrogradely labelled neurons, as well as analysis of marker expression in the amygdala were completed with a PANORAMIC MIDI II (20x (NA 0.8); 3DHistech, Hungary) device and the manufacturer’s official software (CaseViewer 2.4, SlideMaster 2.5) for every 6th slice (i.e., at 300 μm resolution).

### Identification and separation of different amygdala subnuclei

In order to delineate the borders of different amygdala subnuclei and ITC clusters (Extended Data Fig. 5-6.) in a biologically relevant manner we developed an immunohistochemically-based segmentation (IHCS) technique using the following molecular markers: choline acetyltransferase (ChAT), vesicular acetylcholine transporter (VAChT), dopamine- and cAMP-regulated phosphoprotein of molecular weight 32 kDa (DARPP32, also known as PPP1R1B), forkhead box protein P2 (FoxP2), protein kinase C-δ (PKC-δ), somatostatin (SOM) calcitonin gene related peptide (CGRP), calretinin (Calr), and cannabinoid receptor 1 (CB1)^32,117–122^. We compared the expression of these molecular markers to the 5^th^ edition of the Mouse Brain is Stereotaxic Coordinates^93^ and to the Allen Mouse Brain Reference Atlas^94^.

All injections sites, projection patterns or electrode placements (see below) within the amygdala were compared to the expression of one or more relevant markers with multi-channel fluorescent imaging to confirm correct positioning/ROI definition. For simplicity, this confirmation is not always shown in figures. When tracer/virus spillage or track labelling reached neighboring subnuclei or other adjacent regions that could affect the results, animals were excluded from analysis.

### Analysis of anatomical data

#### Axon density analysis

In order to quantify the density of thalamic and cortical axon-arborization in the amygdala, we used high-magnification (63x) confocal Z-stacks (step size: 0.27 μm) from all amygdala subnuclei, at 300 μm antero-posterior (AP) resolution. We aimed to capture stacks where axon density was visibly the highest at each AP levels in each region.

We analyzed the confocal stacks using a custom-made automatic ImageJ macro^16,23,116^ (available at: https://github.com/baabek/Axon-density-analyzer-ImageJ-script.git.). The macro calculated the total axon length (μm) for each stack and channel and the total axon lengths were summated for each subnuclei in each animal. Then, the total axon lengths were compared to the total stack volume (ROI area size (µm^2^) * number of slices * step size (µm) = total stack volume (μm³)) for each stack to calculate the relative axon density (∑ axon length (µm)/∑ stack volume (μm³)). Then relative axon densities were averaged for each subnuclei and each innervation source per animal.

#### Retrograde distribution and intra-amygdala connectivity analysis

To quantify the number of thalamic neurons projecting to different amygdala subnuclei, we used brightfield images at 20x magnification. Labelled neurons were manually registered in each coronal section at 300 μm AP resolution with the built-in tool of CaseViewer 2.4 (3DHistech, Hungary). Only neurons clearly distinguishable from the background were registered. Weakly labelled or ambiguous neurons were excluded from the analysis. Thalamic regions were identified by using anatomically matched calretinin-stained sections^16,23^. The total number of labelled thalamic cells, as well as the number of cells in different thalamic nuclei were then registered per amygdala subnuclei per animal.

For intra-amygdala connectivity analysis, brightfield images (20x magnification) were matched with the corresponding fluorescent images stained with amygdala subregion-specific markers. During analysis, borders of amygdala subnuclei were registered for each plane in ImageJ (NIH) using the default selections tool. All labelled neurons were then registered manually. Only neurons clearly distinguishable from the background were registered. Weakly labelled or ambiguous neurons were excluded from the analysis. The area of each subnuclei (mm^2^) was calculated using the built-in measurement function of ImageJ. Finally, a relative density of neurons (cells/mm^2^) was calculated for each intra-amygdala pathway in each animal. Retrograde data was qualitatively confirmed with classical (5% BDA) anterograde tracing (for LA, BLA, BLP, BMA and BMP) and CAV2-Cre mediated viral tracing (for LA and BLA).

Whole-brain level anterograde connections outside the amygdala were investigated in a qualitative manner using classical (5% BDA) anterograde- (for LA, BLA), Cre-dependent- (LT, dMT, BLA) and CAV2-Cre mediated viral tracing (for LA, BLA) at 300 μm AP resolution. Both fluorescent and brightfield images (see above) were used to reveal anterograde connections of these brain regions.

#### Thalamo-amygdala retrograde tracing

A total of 30 animals were included (*n* = 4: LA, BLA, BMA, Astr, CeL/C, CeM; *n* = 3: BLP, BMP). Cell counts were averaged per amygdala subnucleus. After confirming the homogenous distribution (Levene’s Test for Equality of Variances) a one-way ANOVA test was applied to reveal significant differences in thalamo-amygdala innervation strengths. Percentage values were also calculated for the same dataset to reveal differences in innervations originating from different thalamic nuclei.

#### Thalamo-amygdala and cortico-amygdala axon densities

Data from a total of 4 (thalamus) or 5 (cortex) animals were included in the analysis. Relative axon density values were calculated in each animal by summing the total registered axon length and dividing by the corresponding total volume for each amygdala subnuclei. Values were then averaged across animals per amygdala subnuclei. At least 2 data points per thalamic source per subnuclei was included in all animals. One-way ANOVA with Tukey’s post-hoc test was applied to reveal significant differences in thalamic innervation between amygdala subnuclei.

### Probabilistic tractography

#### Human sample

A dataset of 113 healthy human participants (mean ± SD = 24.5 ± 4.33 years; 65 females) was used for the tractography analyses. All participants were right-handed and had normal or corrected-to-normal vision. No participant had a history of major medical, neurologic, or psychiatric disorders. The study protocol was approved by the Ethics Committee of the Basque Center on Cognition, Brain and Language and was conducted in accordance with the Code of Ethics of the World Medical Association (Declaration of Helsinki) for experiments involving human participants. Before their inclusion in the study, all participants provided informed written consent. Participants received monetary compensation for their participation.

#### Data acquisition

Whole-brain MRI data acquisition was conducted on a 3 T whole-body MRI scanner (Prisma Fit, Siemens Medical Solutions) using a 64-channel whole-head coil. The MRI acquisitions included one T1-weighted (T1w) structural image and diffusion-weighted imaging (DWI) sequences. High-resolution MPRAGE T1-weighted structural images were collected with the following parameters: TR = 2530 ms; TE = 2.36 ms; flip angle = 7°; field of view = 256 mm; voxel size = 1 mm isotropic; 176 slices. In total, 100 diffusion-weighted images were acquired with an anterior-to-posterior phase-encoding direction and 50 isotropically distributed diffusion-encoding gradient directions. The 100 diffusion-weighted images included 50 images with a b-value of 1000 s/mm^2^ and 50 images with a b-value of 2000 s/mm^2^. Twelve images with no diffusion weighting (b-value = 0 s/mm^2^) were obtained for motion correction and geometrical distortion correction, which comprised five images with the same phase-encoding direction as the DWI images and seven images with a reversed-phase encoding direction (posterior to anterior). Both DWI and b0 images shared the following parameters: TR = 3600 ms; TE = 73 ms; flip angle = 78°; voxel size = 2 isotropic; 72 slices with no gap and a multiband acceleration factor of 3.

#### Tractography

The white-matter pathways between MD seeds and PFC were estimated using the Reproducible Tract Profiles 2 (RTP2) pipeline^123^. The T1-weighted image was used to acquire ROIs and to register the DWI to individual space. The amygdala and thalamus were defined in individual T1 space using tools implemented in FreeSurfer (http://surfer.nmr.mgh.harvard.edu/) and thalamic segmentation^124^. Temporal and frontal cortical regions were defined by the HCP atlas^125^ and transformed from MNI space into individual space using a nonlinear transformation in Advanced Normalization Tools (ANTs; http://stnava.github.io/ANTs).

The DWI data were preprocessed using MRtrix functions^126^ in the following steps: (1) data denoising based on random matrix theory, which exploits data redundancy in the patch-level principal component analysis domain using *dwidenoise*^127^; (2) Gibbs Ringing correction using *mrdegibbs*^128^; (3) susceptibility-induced distortions and motion correction with the FSL topup and eddy tools called by *dwifslpreproc*^129^; (4) B1 field inhomogeneity correction with *dwibiascorrect* and Rician background noise removal with *mrcalc* and lastly (5) a rigid transformation matrix to align the DWI images to the corresponding T1w image using ANTs.

White-matter pathways were reconstructed from the preprocessed DWI data. We first modeled the diffusion information at the voxel level to obtain a map of preferred directions with fiber orientation distributions (FODs). For this modeling, we used the MRtrix3 CSD algorithm^130^, as it can discern crossing fibers and provide more than one direction in each voxel. Next, streamline tracking was performed separately for pathways between (1) lateral thalamus and amygdala (LT-Amy), (2) medial thalamus and amygdala, (3) lateral thalamus and temporal cortex (LT-TeCx), (4) medial thalamus and prefrontal cortex (MT-PFC), (5) temporal cortex and amygdala (TeCx-Amy), (6) prefrontal cortex and amygdala (PFC-Amy). A full list of individual pathways and groups can be found in *Source Data Table*. The tracking was performed on the estimated FODs using the MRtrix iFOD2 algorithm^131^ with the following parameters: step size, 1 mm; FODs amplitude threshold, 0.1; angle threshold, 45°; maximum length, 200 mm; minimum length, 10 mm.

Tract streamline counts were then summed for each main stream and averaged across all 113 participants. Since a test for Equality of Variances (Leven’s) revealed a non-homogeneous distribution, we used Kruskal-Wallis non-parametric test with Dunn’s post-hoc comparisons to examine statistically significant differences between main stream tract counts.

### In vivo electrophysiology

In vivo recordings were performed 4–12 weeks after viral injections under urethane anesthesia (20 m/m % dilution; 0.005ml/1g). The head of the animal was fixed in a stereotaxic frame. Neuronal activity from the AMY (AP –1.5 to -2, ML 2.8 to 3.6, DV 4.5 mm) was monitored with high density silicon probes (Buzsáki 64/256; Neuronexus), stained with fluorescent DiI. All recordings were done ipsilaterally to the sites of viral injection. Wideband neural data (0.1-7500Hz) were high-pass filtered (0.3 Hz), amplified (400 V/V) by a 256-channel amplifier, and digitized at 20 kHz (RHD2000, Intan Technologies and KJU-1001, Ampliplex). A screw driven into the occipital bone served as reference electrode. For optogenetic activation, the optic fiber was placed above the amygdala, where the LT-Calr+ (*n* = 5 animals) or dMT-Calr+ (*n* = 3 animals) fibers were activated with blue light laser pulses (Laserglow Technologies). Sensory stimulus was delivered as footshocks via bipolar electric stimulation on the left rear feet (100 ms, 0.1 mA). Optogenetic activations were done with multimode optic fibers (105 μm core diameter) and consisted of 473 nm blue light pulses. With a single cue type, low-intensity optogenetic activations (5 ms, 1 mW) were repeated at 0.1 Hz for 3x10 minutes with 5-minute interblock intervals; 6x10 foot shock stimuli (100 ms, 1 mA) were delivered with 30 s interstimulus intervals and a few minutes interblock intervals. These recordings were done in the Oscillatory Neuronal Networks Research Group at the University of Szeged (group leader: Dr. Antal Berényi). The lasers and the current generator (Multi Channel Systems) were triggered by analog signals delivered with a National Instruments acquisition board (USB-6353). Analog trigger pulses were also delivered to the evaluation board and registered in parallel with neural data for synchronization purposes. All stimulation protocols were executed with custom-written MATLAB (Mathworks) codes. After recordings, animals were transcardially perfused (see above).

### Neural data processing

Spike detection and automatic clustering were performed using Kilosort2.5^132^. Afterwards, cell grouping was manually refined: a group of spikes was considered to be generated by a single neuron if they formed a discrete, well-isolated cluster and had an autocorrelogram with a clean refractory period (if average bin values of the first 2 ms did not reach the autocorrelogram’s asymptote line). Putative cell grouping and further data analyses were carried out using CellExplorer^45^ and basic, custom-built scripts in MATLAB and Python for plotting. All the presented electrophysiological data are derived from individually identified single units (clustered cells). Clustered cells were separated into 4 different putative neuronal subtypes, pyramidal cells (PC, *n* = 633), narrow Interneurons (NI, *n* = 56), wide interneurons (WI, *n* = 92), and medium spiny neurons (MSN, *n* = 126), based on firing rate (FR), autocorrelogram shape (ACG Fit) and trough-to-peak value. Location of a given putative cell-type within a given AMY subnucleus was selected based on the position of the recording site with the highest amplitude spike. Location of recording sites was verified by using the DiI-labeled trace for a virtual anatomical reconstruction of the electrode tracks using our IHCS mapping. The used brain atlas coordinates for AMY and the electrode configurations allowed us to record simultaneous extracellular neuronal signals (spikes) from most of the IHCS identified amygdala subnuclei with some limitations. Having a spatial resolution of ∼100 µm ^133^, spiking activity in the capsular part of the central amygdala (CeC) could not be reliably differentiated from those originated in the lateral part of the central amygdala (CeL). Similar scenario was present in case of spiking activity in the anterior (BMA) and posterior parts of the basomedial amygdala (BMP). Thus, in our analysis, we grouped them together and named them as CeL/C and BMA/P.

Z-score values were calculated as PSTH either pooled from all putative cells within a given subnucleus (Fig. 2i and Extended Data Fig. 13a), or for individual putative cell types (in case of the proportional distribution of individual cell latency, Extended Data Fig. 13b) with regards to -800 ms baseline period (calculated from the onset of each stimulus). Latency of individual cell types were calculated as the 3rd consecutive significant bin (1 bin = 1 ms, *z* > 1.93, *P* > 0.05). The latency of subnucleus-specific activation (Fig. 3a) was determined as positive crossings of the mean + SD cumulative sum curve calculated from pooled post-stim spiking activity (0-30 ms) above a threshold of significance. The threshold of significance was determined as mean + 3*SD cumulative sum of baseline activity (-30 to 0 ms, pre-stimulus). The activation probability of a given amygdala subnucleus (Fig. 3b) was calculated as the number of activated putative neurons which had a latency value within the given time bin, divided by the sum of the same neurons and neurons that were not activated (i.e. the ones without measurable latency and significant activation in any of the examined time bins).

### Correlation between anatomical and electrophysiological data

To examine the strength of the thalamo-amygdala or intra-amygdala connectivity, we correlated our anatomical and electrophysiological datasets. In case of the thalamo-amygdala connectivity, the thalamo-amygdala axon density values (see above) were normalized (x_normalized_ = (x - x_minimum_) / range of x) per thalamic source. Normalized values were then averaged across animals per amygdala subnucleus per thalamic source. Correlations between these values derived from anatomy and electrophysiology were then calculated using one-tailed Pearson’s correlation test for positive correlations.

In case of the intra-amygdala connectivity, we used the dataset of retrogradely labelled intra-amygdala neurons, quantified as described above for a total of 27 animals (*n* = 4: LA, BLA, Astr; *n* = 3: BLP, BMA, BMP, CeL/C, CeM). Density (cells/mm^2^) and the values were normalized for the whole dataset (x_normalized_ = (x - x_minimum_) / range of x). Normalized values were then averaged across animals for amygdala subnucleus pairs to generate “connection strength” values for each intra-amygdala pathway. For simplicity, data in figures is presented as “output” for each subnucleus. Correlations between these values were then calculated using one-tailed Pearson’s correlation test for positive correlations.

For electrophysiological data, Z-score values were calculated from PSTH for each putative cell type with regards to -800 ms baseline period (calculated from the onset of each stimulus) and averaged across animals per amygdala subnucleus within consecutive post-stimulus intervals to group responses into short-latency (putative thalamo-amygdala) and long-latency (putative intra-amygdala) responses (short: 5-9 ms, long: 10-14 for LT/dMT and short:15-19 ms, long: 20-24 ms for sensory stimulation).

### Statistical analysis

All animals were randomly assigned to experiments with no regards to their sex. No human participants were excluded from analysis even if tract counts were 0 for 1 or more individual tracts. We used JASP (Version 0.95.2; JASP Team (2025)) for statistical analysis. Raw data and detailed statistics can be found in Source Data Tables and Supplementary Tables arranged according to corresponding figures.

## Author contributions

Á.B and F.M. conceived the initial concept of the study; Á.B., K.K., F.C., P.M.P-A. and F.M conceptualized and designed the study; K.K along with P.B., and A.M. performed the electrophysiological recordings; P.B. K.K. and S.B. analyzed all recordings; A.B. and F.M. supervised the electrophysiological recordings; Á.B., A.V.B., J.B. and F.M. acquired the anatomical data; Á.B., B.B. and F.M. analysed the anatomical data; L.M. acquired the human tractography data; F.C. and P.M.P-A. supervised the human tractography; B.B. provided statistical assistance; A.B. provided resources; Á.B., B.P. and F.M. wrote the original draft of the paper. All authors contributed to writing, reviewing and editing.

## Acknowledgements

We thank Z.J. Huang, Balázs Rózsa and Norbert Hájos for providing us transgenic mice; Tamás Herczeg, Réka Erdős, Anna Fehér and Katalin Varga for laboratory assistance; László Acsády, István Katona and Norbert Hájos for sharing antibodies. We thank Andrew Holmes for comments and discussions about the manuscript. We also thank the Institute of Molecular Life Sciences of the HUN-REN Research Centre for Natural Sciences for the use of Zeiss microscope, the Light Microscopy Center and the Virus Technology Unit of HUN-REN Institute of Experimental Medicine for their help. This work was supported by the following funding agencies: Ministry of Innovation and Technology of Hungary from the National Research, Development and Innovation Fund (NKFI-138836, NKFI-153312 and KKP126998 to F.M.; NKFI-135285, NKFI-144583 to S.B.; NKFI-124034 to B.B.; HU-RIZONT-2024-00003 to A.M.); Cooperative Doctoral Program (KDP-2020-1015461 to A.M.); Hungarian Brain Research Program (2017-1.2.1-NKP-2017-00002 to F.M.); New National Excellence Program of the Ministry for Innovation and Technology (ÚNKP-21-5-ÁTE-2 to F.M.; ÚNKP-21-3-II-BME-61, ÚNKP-20-3-II-BME-24 and ÚNKP-18-3-II-BME-55 to Á.B; ÚNKP-19-3-III-PPKE-68 to K.K.; ÚNKP-22-2-III-ELTE-539 to A.V.B., ELKH SA-48/2021); New National Excellence Program of the Ministry for Culture and Innovation from the source of the National Research, Development and Innovation Fund (EKÖP-2024-87 to P.B., EKÖP-2024-276 to A.V.B., 2025-2.1.1-EKÖP-2025-00014 to A.M.). P.M.P-A. was supported by a grant from the Spanish Ministry of Science and Innovation (PID2024-159163NB-I00), and acknowledges funding to the BCBL from the Basque Government through the BERC 2022–2025 program and from the Spanish State Research Agency through the Severo Ochoa Center of Excellence accreditation CEX2020-001010-S). FM was a János Bolyai Research Fellow.

---

**Extended Data Figure 1:**
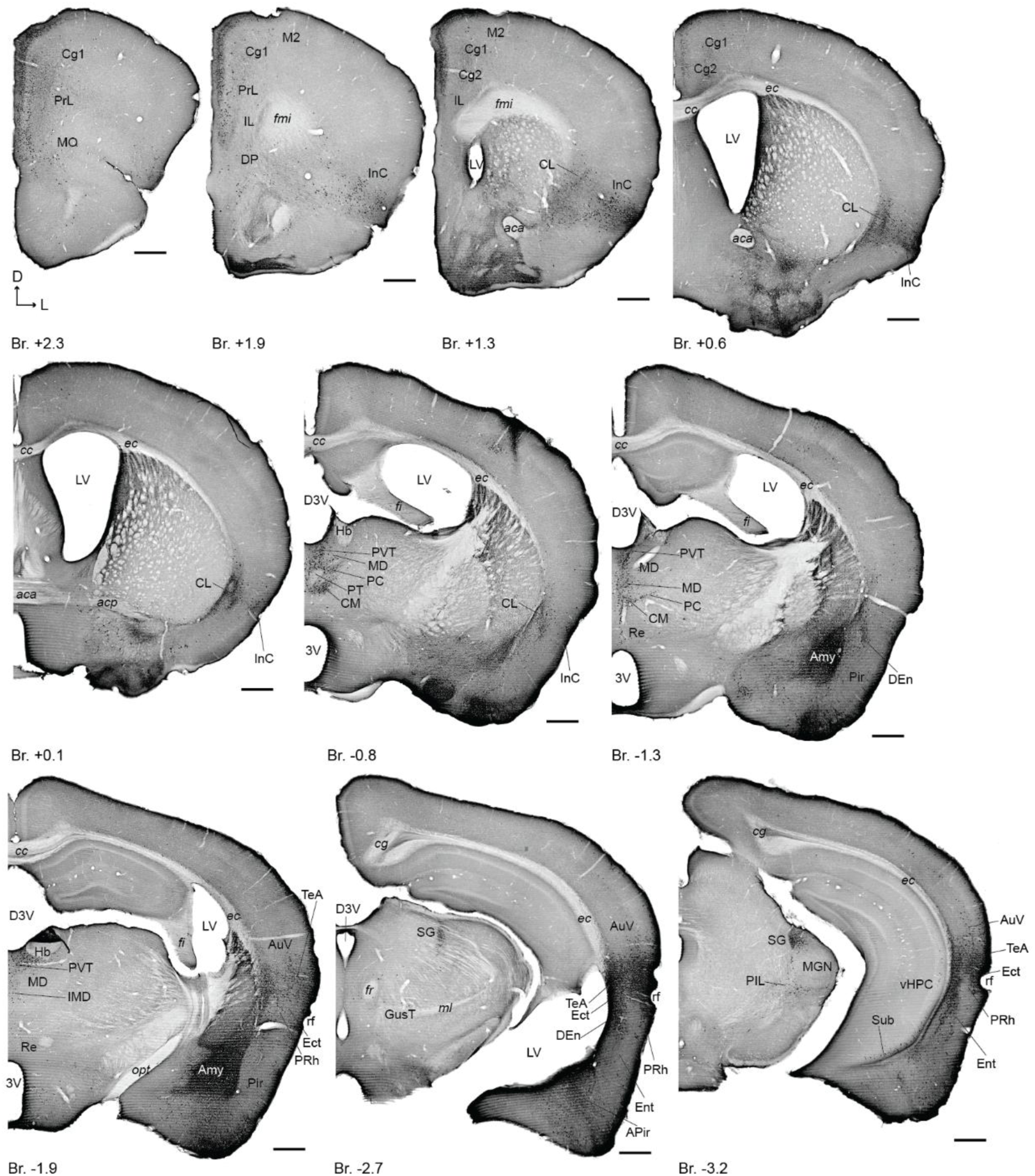
Brain-wide distribution of the retrogradely-labeled neurons from the amygdala (related to Fig. 1). For experimental design see Fig. 1a. Images taken from the same animal as in Fig. 1b-c. Injection site in the amygdala is indicated in white font. Scale bars, 500 µm. For abbreviations see *Supplementary Tables*.

**Extended Data Figure 2:**
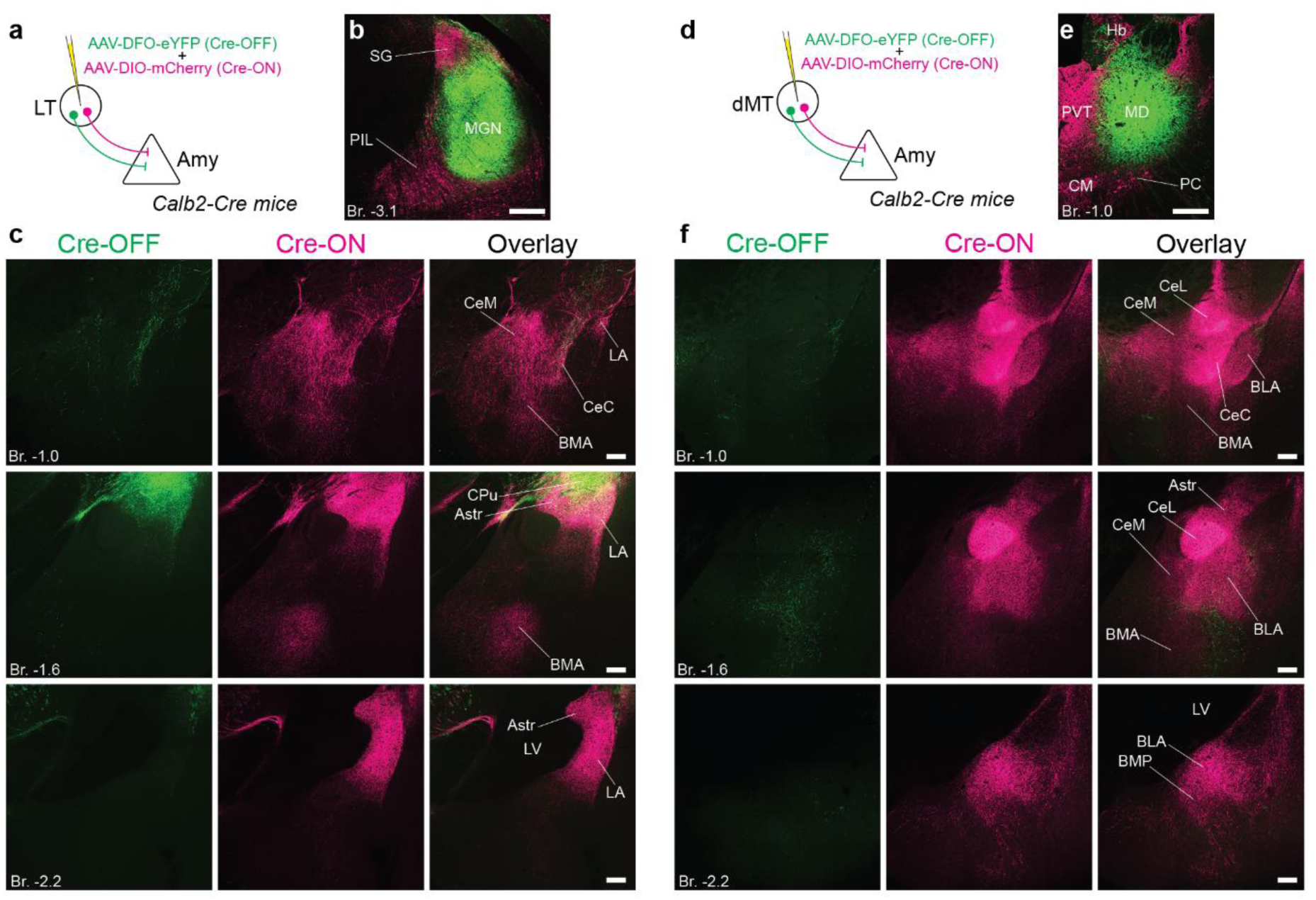
Calretinin-positive cell-type specific origin of the thalamo-amygdala networks (related to Fig. 1). **a**, **d**, Experimental designs for Calr-positive (Cre-ON) and -negative (Cre-OFF) anterograde viral tracing in the LT (**a**) and dMT (**d**). **b**,**e**, Double AAV (Cre-ON – magenta, Cre-OFF – green) injection site in the LT (**b**) and dMT (**e**). **c**,**f**, Distribution of Calr-positive (magenta) and Calr-negative (green) LT (**c**) and dMT (**f**) axons in the amygdala at three different AP levels. Scale bars, 200 µm.

**Extended Data Figure 3:**
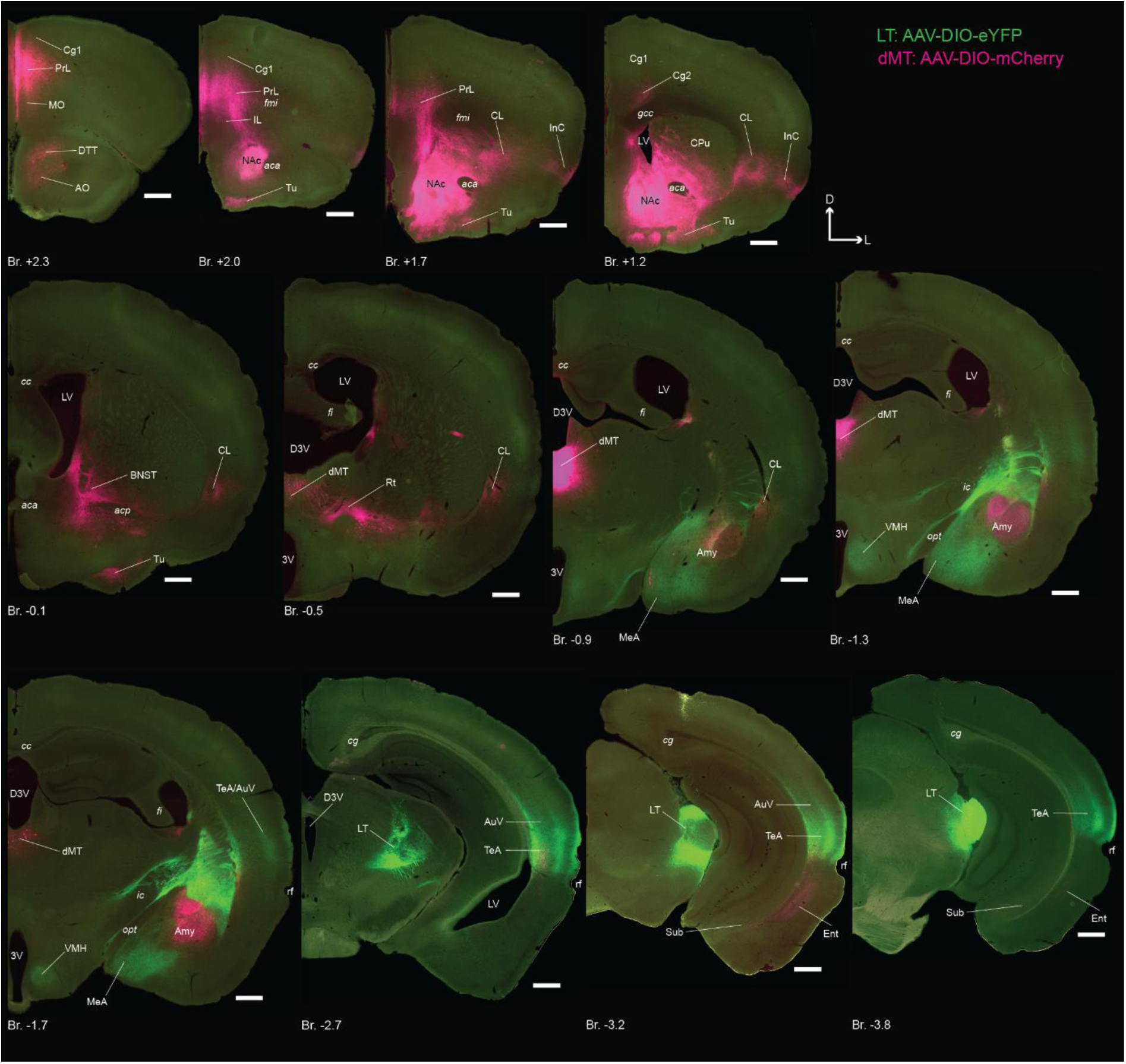
Whole-brain distribution of LT and dMT axonal arbors (related to Fig. 1). For experimental design see Fig. 1d. Images taken from the same animal as in Fig. 1e. Images at Br. +2.0, -1.7 and -3.2 are also shown in Fig 1e. Scale bars, 500 µm. For abbreviations see *Supplementary Tables*.

**Extended Data Figure 4:**
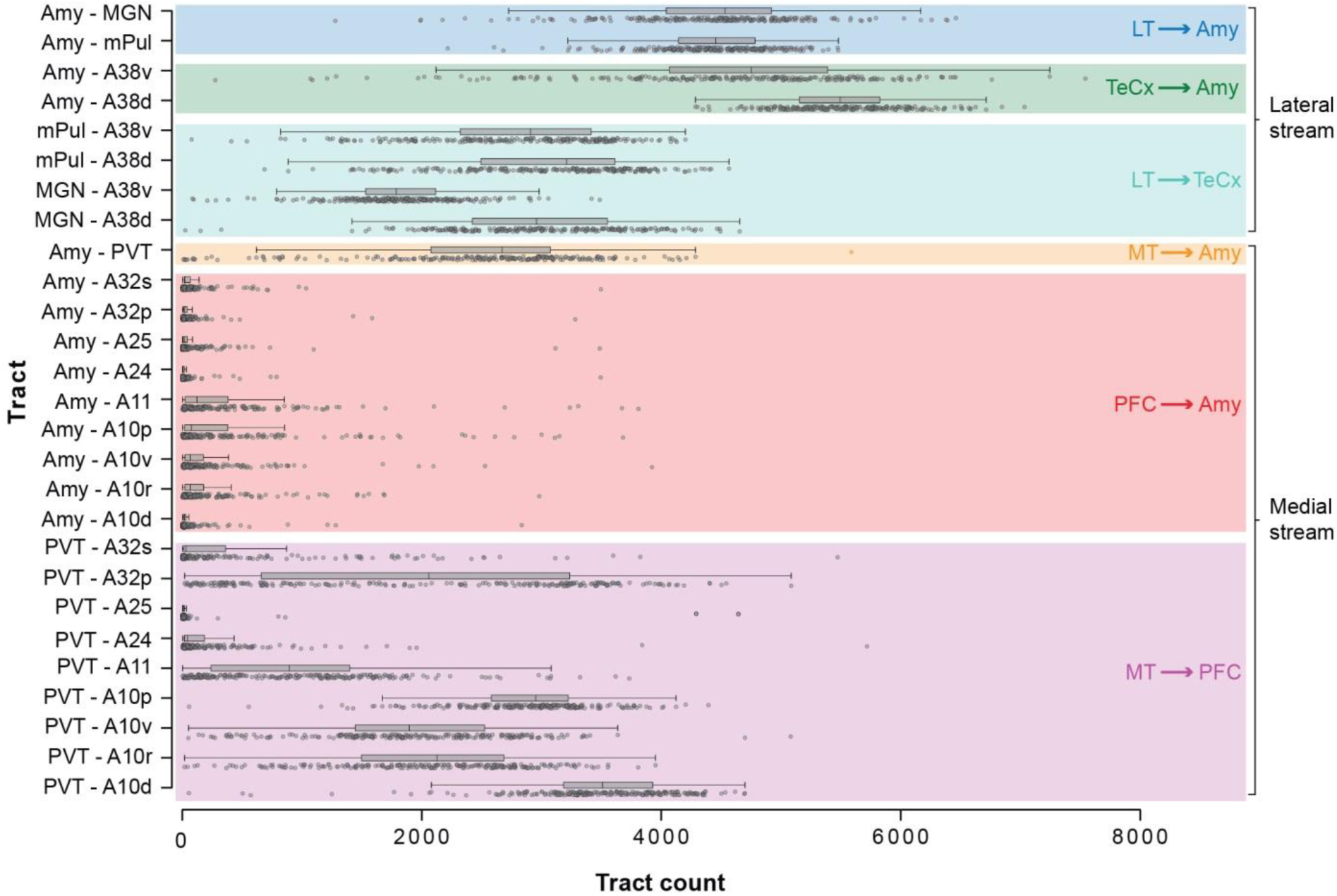
Individual cortical and thalamic tracts of the lateral and medial streams connected to the amygdala in humans (related to Fig. 1). Box-plots indicate the interquartile range and distribution density and the whiskers show 1.5x interquartile ranges. Circles indicate data from individual patients. Same dataset as in Fig 1f-i. For details and abbreviations see *Source Data Tables* and *Supplementary Tables*.

**Extended Data Figure 5:**
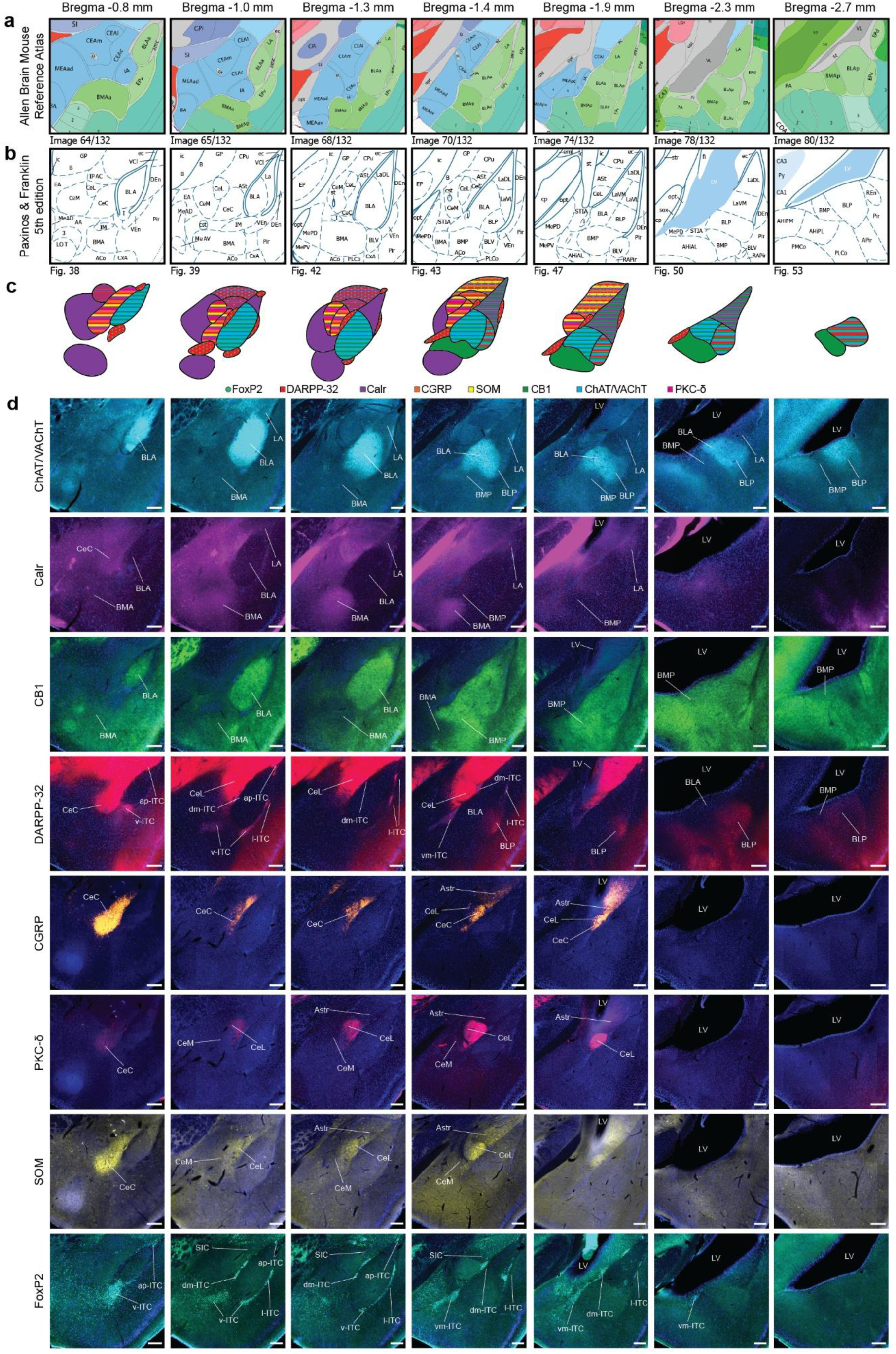
Immunohistochemically-based segmentation (IHCS) for reliable delineation of individual amygdala subnuclei (related to Fig. 2). **a,** Amygdala parcellation according to the Allen Mouse Brain Reference Atlas (Allen Reference Atlas – Mouse Brain, n.d.)(Allen Reference Atlas – Mouse Brain, n.d.)(Allen Reference Atlas – Mouse Brain, n.d.) in coronal sections at 7 different AP levels. **b,** Amygdala parcellation according to Paxinos and Franklin’s The Mouse Brain Atlas (5^th^ edition) (Paxinos and Franklin, 2019)(Paxinos and Franklin, 2019)(Paxinos and Franklin, 2019)(Paxinos and Franklin, 2019)(Paxinos and Franklin, 2019)(Paxinos and Franklin, 2019)(Paxinos and Franklin, 2019)(Paxinos and Franklin, 2019)(Paxinos and Franklin, 2019) at similar AP levels as in (**a**). **c,** Amygdala parcellation used in this article based on molecular marker expressions at similar AP levels as in (**a, b**). Overlapping expression is represented with stripes. Nucleus-selective marker (FoxP2) is represented with dots. **d,** Representative fluorescent images showing the expression of 8 different molecular markers (ChAT/VAChT – cyan, Calr – purple, CB1 – green, DARPP-32 – red, CGRP – orange, PKC-δ – magenta, SOM – yellow, FoxP2 – turquoise) used in this article to define amygdala subnuclei at similar AP levels as in (**a-c**). Scale bars, 200 µm. For abbreviations see *Supplementary Tables*.

**Extended Data Figure 6:**
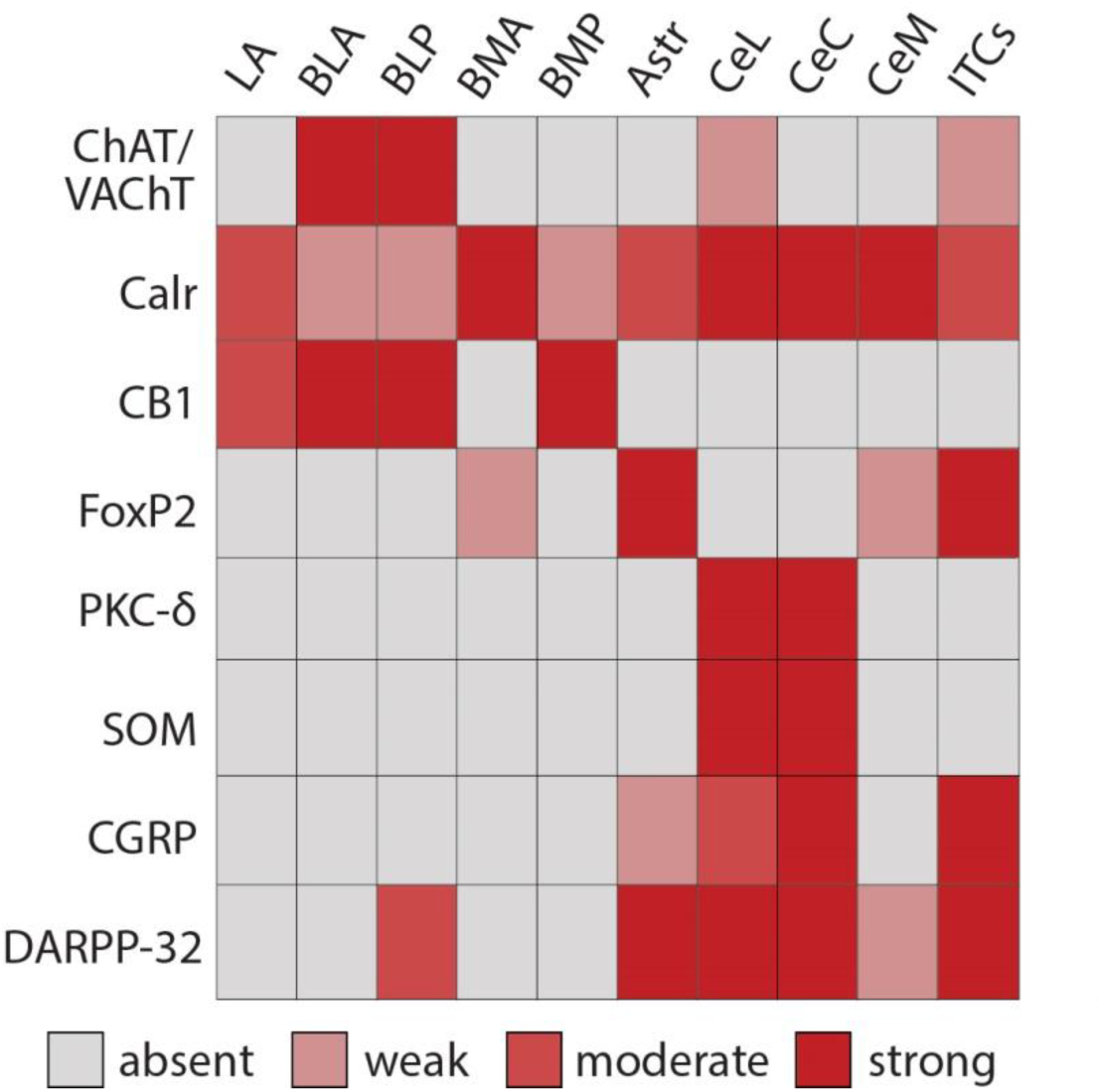
Heatmap summarizing IHCS marker distribution in the amygdala (related to Fig. 2). Darker shades indicate higher expression level based on semi-quantitative analysis (for details and abbreviations see *Supplementary Tables*.)

**Extended Data Figure 7:**
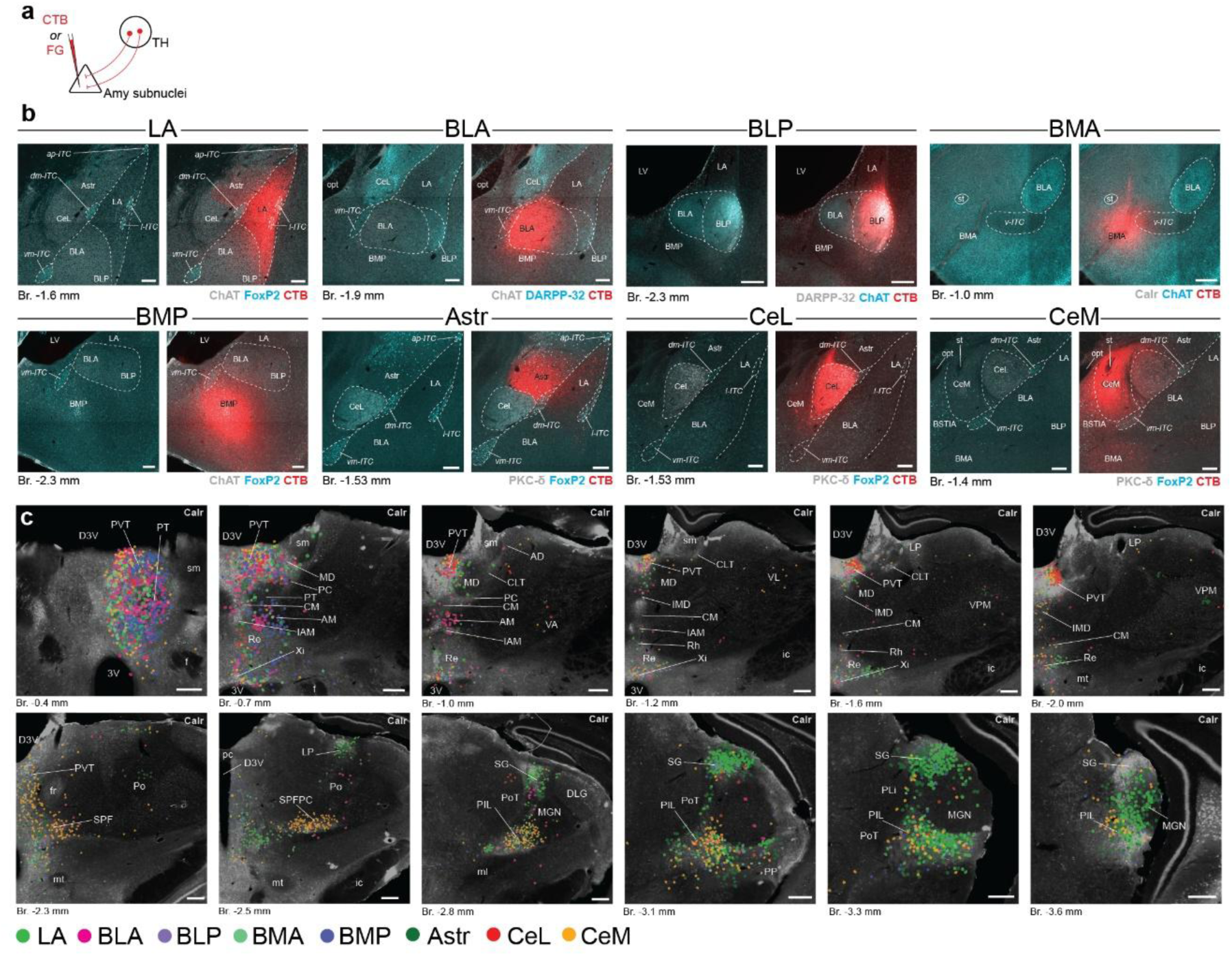
Distribution of LT and dMT neurons projecting to different amygdala subnuclei (related to Fig. 2) **a,** Experimental design for amygdala subnucleus-specific retrograde tracing in the thalamo-amygdala network. **b,** Representative retrograde tracer (red) injection sites in different amygdala subnuclei, aligned with two relevant molecular markers (cyan, grey). Scale bars, 200 µm. **c**, Distribution of amygdala-innervating neurons in the thalamus at 12 AP levels compared to Calr-expression (greyscale). Each dot represents one labelled neuron after individual amygdala subnuclei injection taken from 1-1 representative mouse. Scale bars, 200 µm. For abbreviations see *Supplementary Tables*.

**Extended Data Figure 8:**
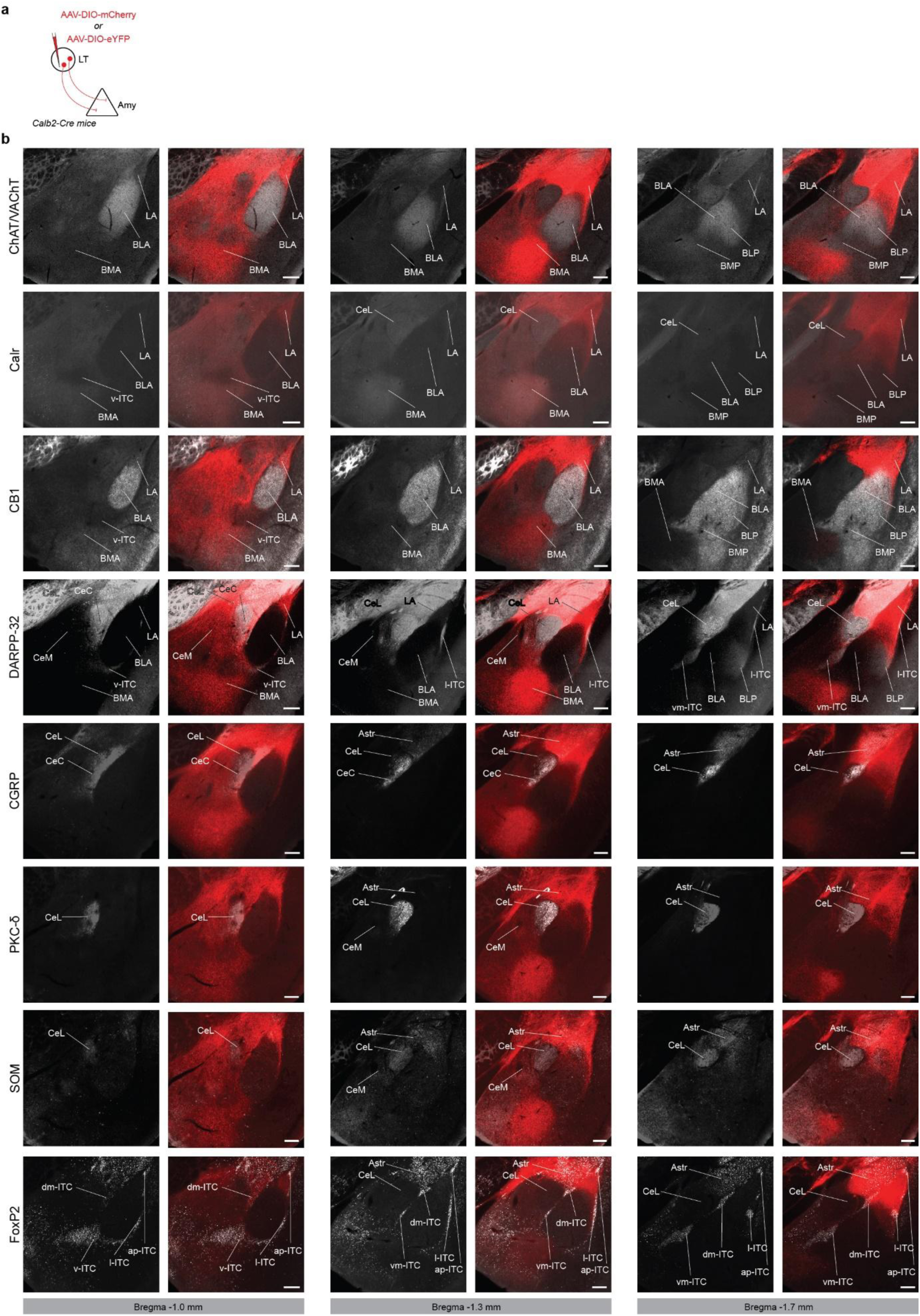
Distribution of LT axons in the amygdala complex according to the IHCS (related to Fig. 2). **a,** Experimental design for AAV-mediated anterograde thalamo-amygdala circuit tracings from LT. **b,** Distribution of AAV labelled LT axons (red) compared to the expression of molecular markers (greyscale) used to define amygdala subnuclei at 3 different AP levels. Note that images with eYFP labeling were recolored to red for uniformity. Scale bars, 200 µm. For abbreviations see *Supplementary Tables*.

**Extended Data Figure 9:**
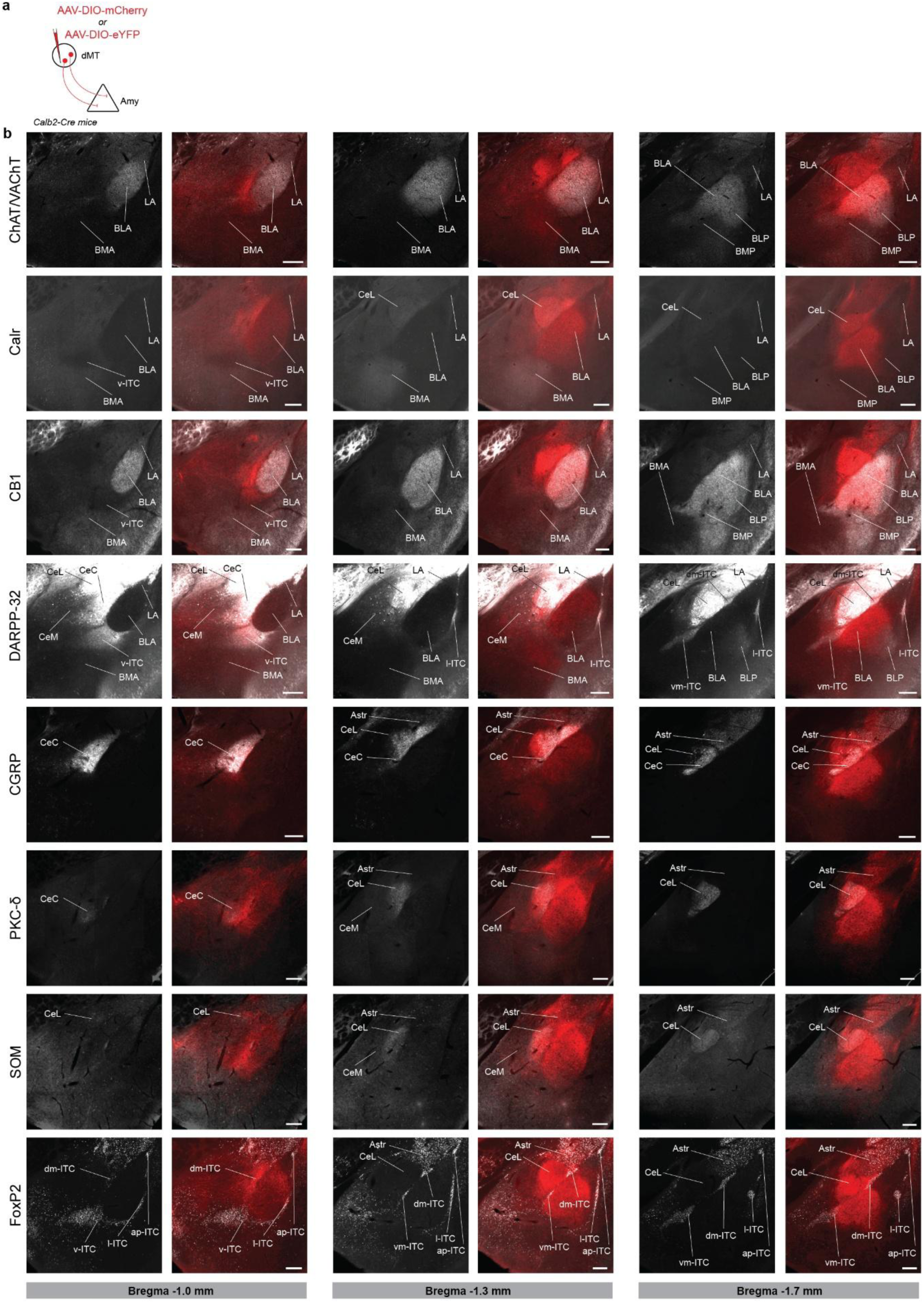
Distribution of dMT axons in the amygdala according to the IHCS (related to Fig. 2) **a,** Experimental design for AAV-mediated anterograde thalamo-amygdala circuit tracings from dMT. **b,** Distribution of AAV labelled dMT axons (red) compared to the expression of molecular markers (greyscale) used to define amygdala subnuclei at 3 different AP levels. Note that images with eYFP labeling were recolored to red for uniformity. Scale bars, 200 µm. For abbreviations see *Supplementary Tables*.

**Extended Data Figure 10:**
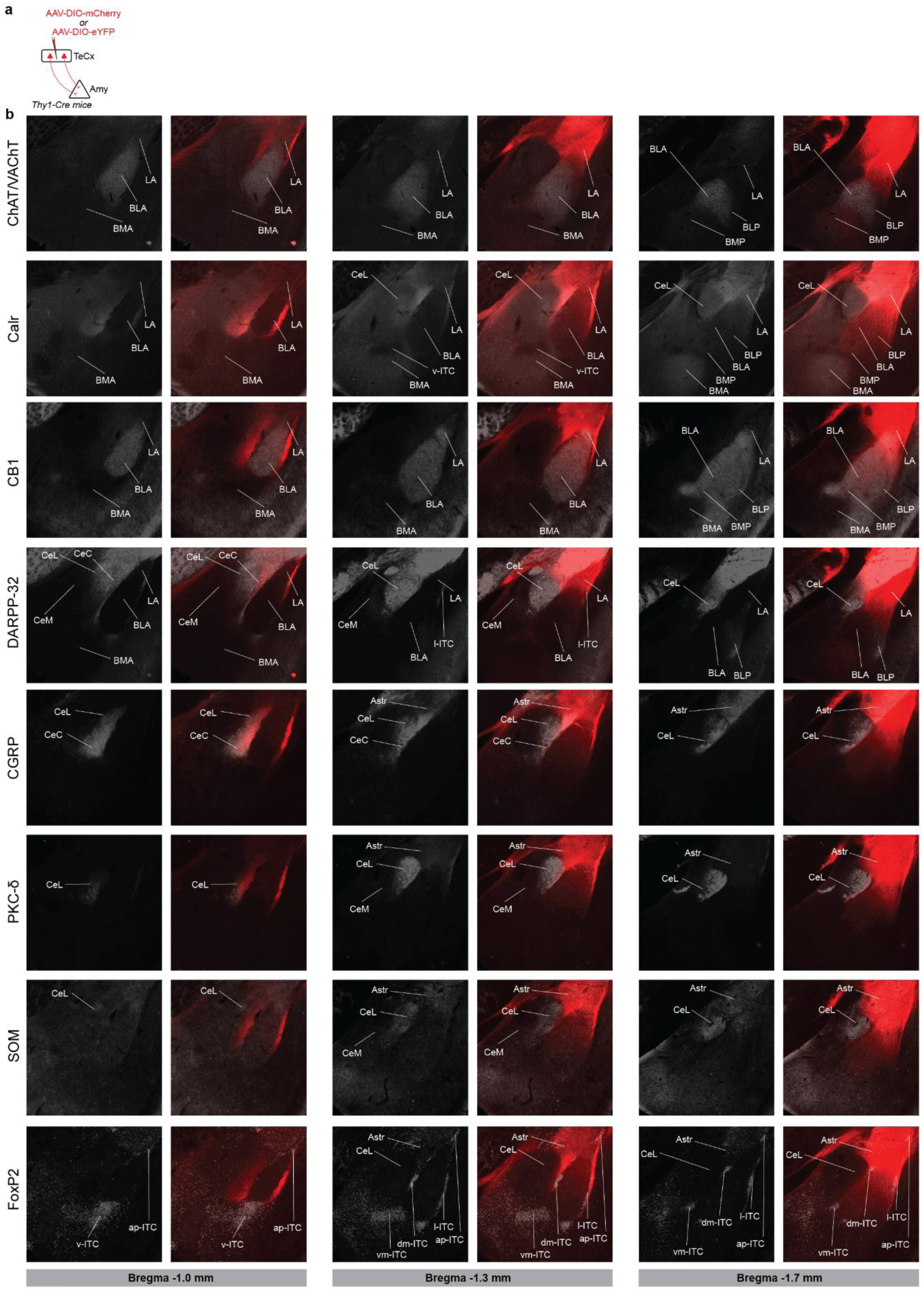
Subnuclear distribution of TeCx axons in the amygdala according to the IHCS (related to Fig. 2) **a,** Experimental design for AAV-mediated anterograde cortico-amygdala circuit tracing from TeCx. **b,** Distribution of AAV labelled TeCx axons (red) compared to the expression of molecular markers (greyscale) used to define amygdala subnuclei at 3 different AP levels. Note that images with eYFP labeling were recolored to red for uniformity. Scale bars, 200 µm. For abbreviations see *Supplementary Tables*.

**Extended Data Figure 11:**
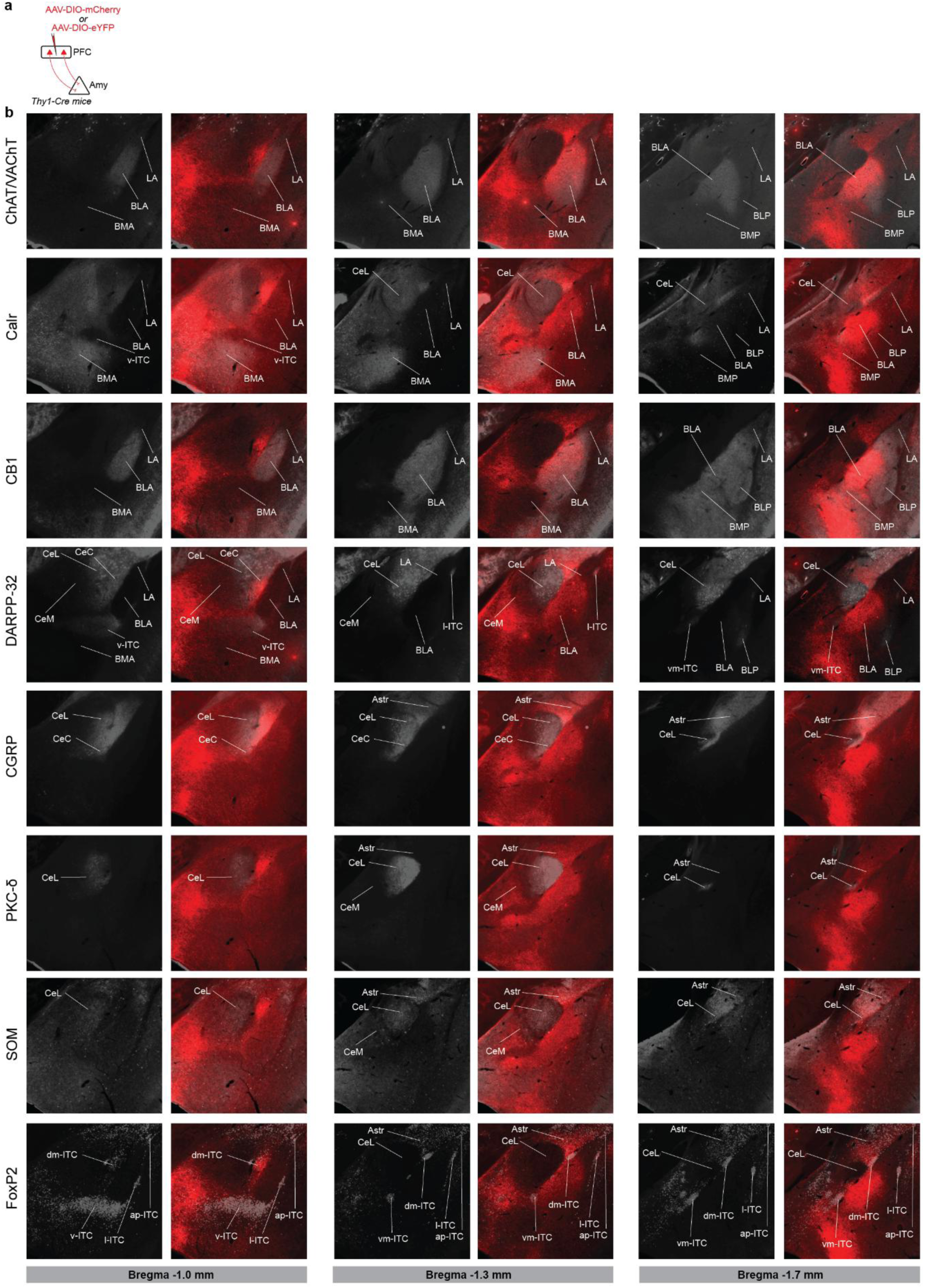
Subnuclear distribution of PFC axons in the amygdala according to the IHCS (related to Fig. 2) **a,** Experimental design for AAV-mediated anterograde cortico-amygdala circuit tracings from PFC. **b,** Distribution of AAV labelled PFC axons (red) compared to the expression of molecular markers (greyscale) used to define amygdala subnuclei at 3 different AP levels. Note that images with eYFP labeling were recolored to red for uniformity. Scale bars, 200 µm. For abbreviations see *Supplementary Tables*.

**Extended Data Figure 12:**
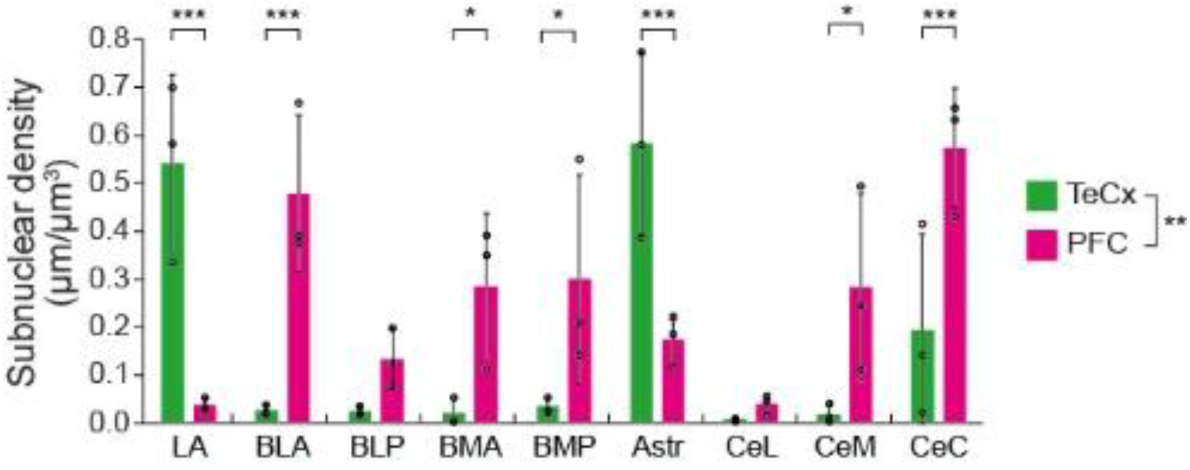
TeCx and PFC innervation of individual amygdala subnuclei (related to Fig. 2 and Extended data figure 10 and 11) Quantification of relative TeCx (green) and PFC (magenta) axon densities at individual amygdala subnucleus level (*n* = 5 mice; one-way ANOVA, Cortex, *F_1,8_* = 8.181, *P* = 0.007, Cortex*Amygdala subnucleus, *F_1,8_* = 11.259, *P* = 8.232*10^-8^, Tukey’s post-hoc test, \**P* < 0.05, \*\*\**P* < 0.001). For detailed statistical data see *Supplementary Tables*.

**Extended Data Figure 13:**
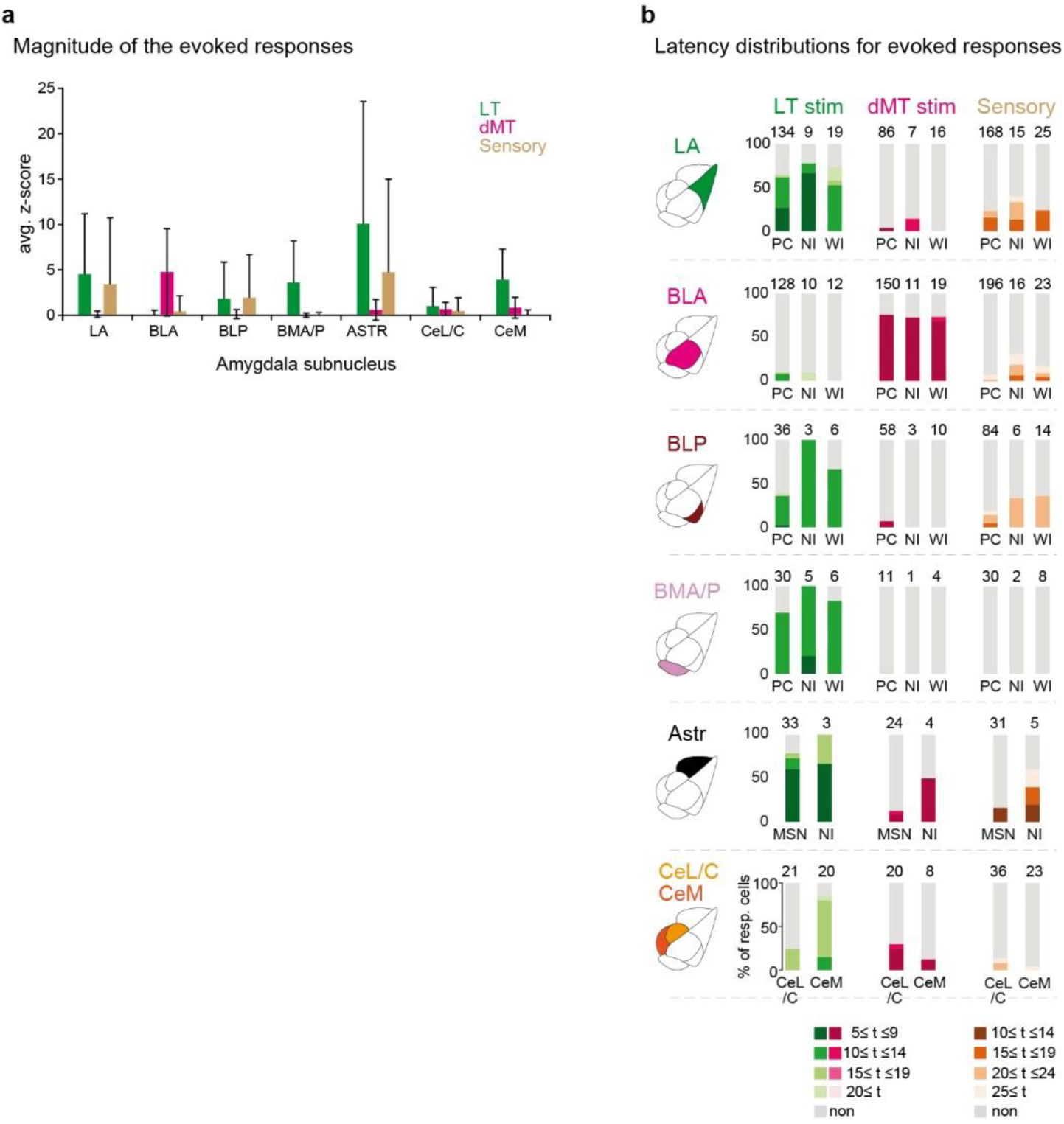
Quantitative analysis for the activation of amygdala neurons evoked by thalamic input and sensory stimulations in relation to their amygdala subnuclear location (related to Fig. 2). **a**, Heterogeneous, amygdala subnucleus-specific responses are evoked by the photostimulation of LT and dMT inputs as well as sensory stimuli. In cases of LT and dMT trials, 0-20 ms time window, while in case of sensory stimulation, 10-30 ms time window was analyzed. (Kruskal-Wallis test; LT, *H(6)* = 148.7*, P =* 1.474*10^-29^; dMT, *H(6) =* 241.1, *P =* 3.077*10^-49^; US, *H(6)* = 30.63, *P* = 2.98*10⁻⁵). Same dataset as in Fig. 3b. **b**, Percentages of responsive neurons with latencies of 5 ms bins within a given amygdala subnuclei in response to LT, dMT or sensory stimuli. Light grey indicates percentages of non-responsive neurons (<100 ms). Numbers above the bars represent the total number of neurons in a given subtype. For detailed statistical data see *Supplementary Tables*.

**Extended Data Figure 14:**
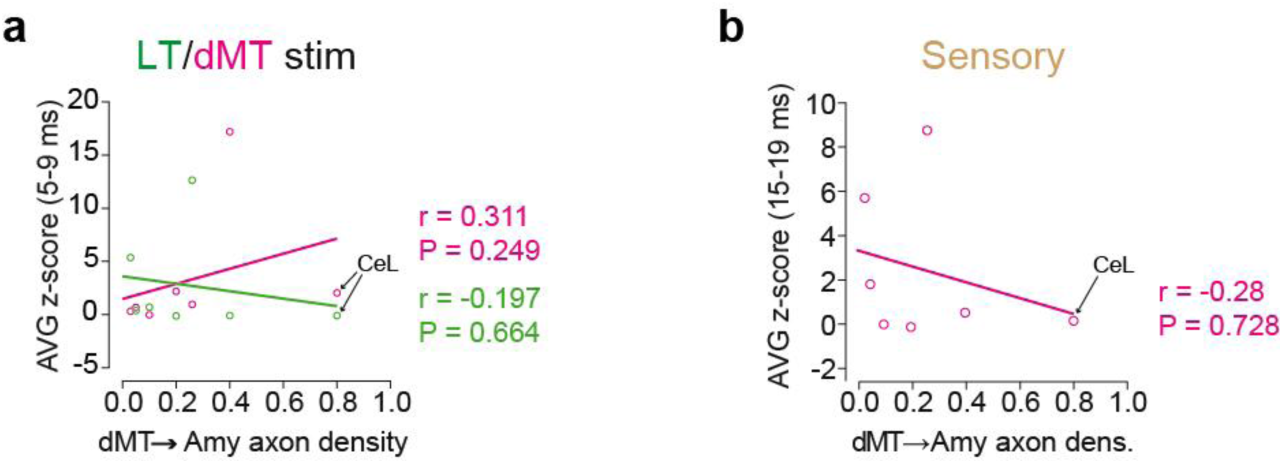
Additional data for the correlations between the thalamo-amygdala anatomical and functional connectivity (related to Fig. 3). **a**,**b**, Same dataset as in Fig 3c,d, but also including CeL/C values. X-axis: normalized axon densities, y-axis: electrophysiological activation obtained with optogenetic and sensory stimulation (analyzed as the averaged z-score calculated in a given post-stimulus interval). Pearson correlation analysis. Circles indicate anatomical and electrophysiological value pairs for each amygdala subnucleus. For detailed statistical data see *Supplementary Tables*.

**Extended Data Figure 15:**
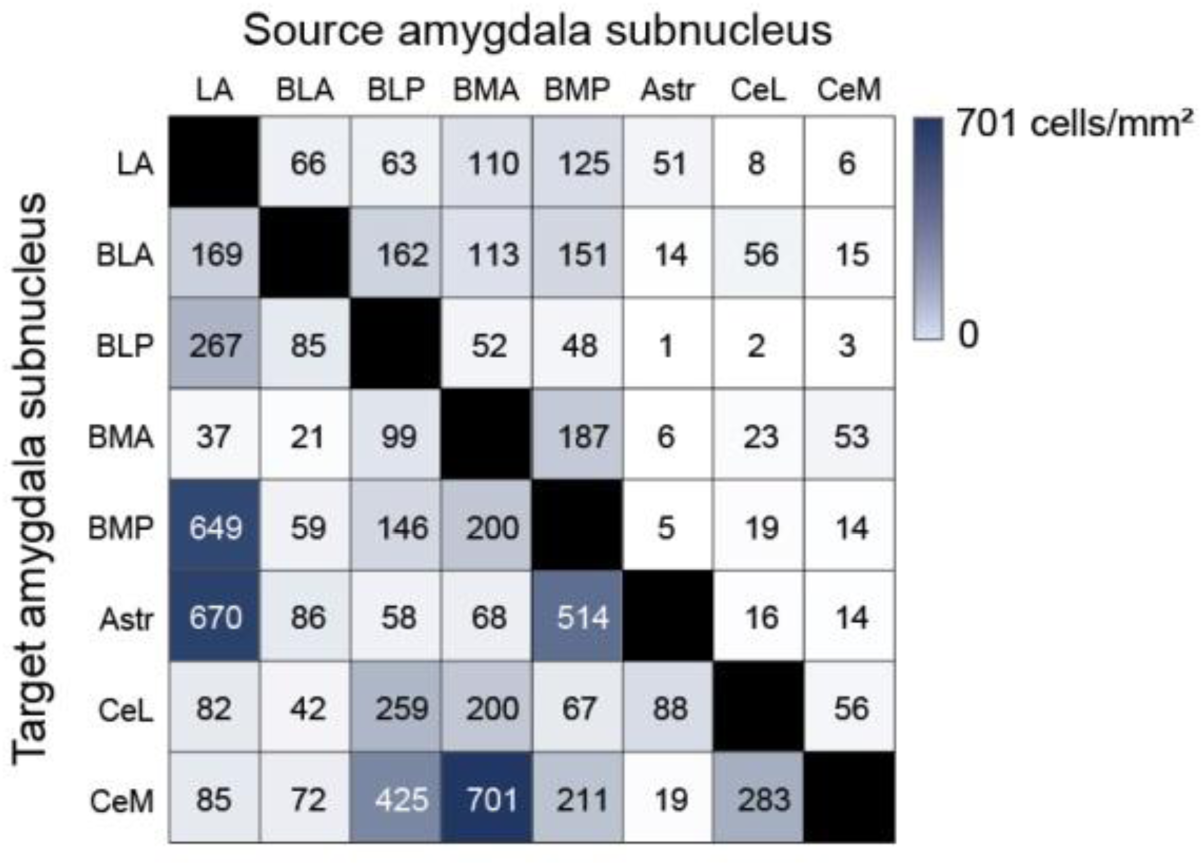
A heatmap summarizing the strength of the intra-amygdala connections (related to Fig. 4). Same dataset as in Fig. 4a-b. Numbers indicate average density (# of cells/mm^2^) values (*n* = 3-4 mice per trial). For details see *Supplementary Tables* and *Source Data Tables*.

**Extended data figure 16:**
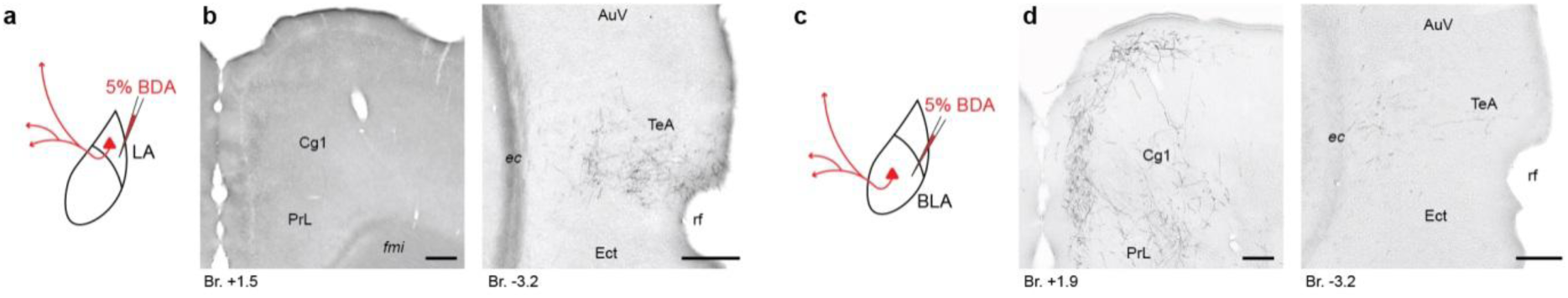
Opposite cortical innervation of LA and BLA neurons (same trials as in Fig. 5). **a**, Experimental design for LA selective anterograde (BDA) tracings in the amygdalo-cortical network in mice. **b**, Representative high magnification images showing weak prefrontal (PrL and Cg1; *left*) and strong temporal (TeA; *right*) cortical axonal projections of LA. **c**, Experimental design for BLA selective anterograde (BDA) tracings in the amygdalo-cortical network in mice. **d**, Representative high magnification images showing strong prefrontal (PrL and Cg1; *left*) and weak temporal (TeA; *right*) cortical axonal projections of BLA. Scale bars, 200 µm. For abbreviations see *Supplementary Tables*.

**Extended data figure 17:**
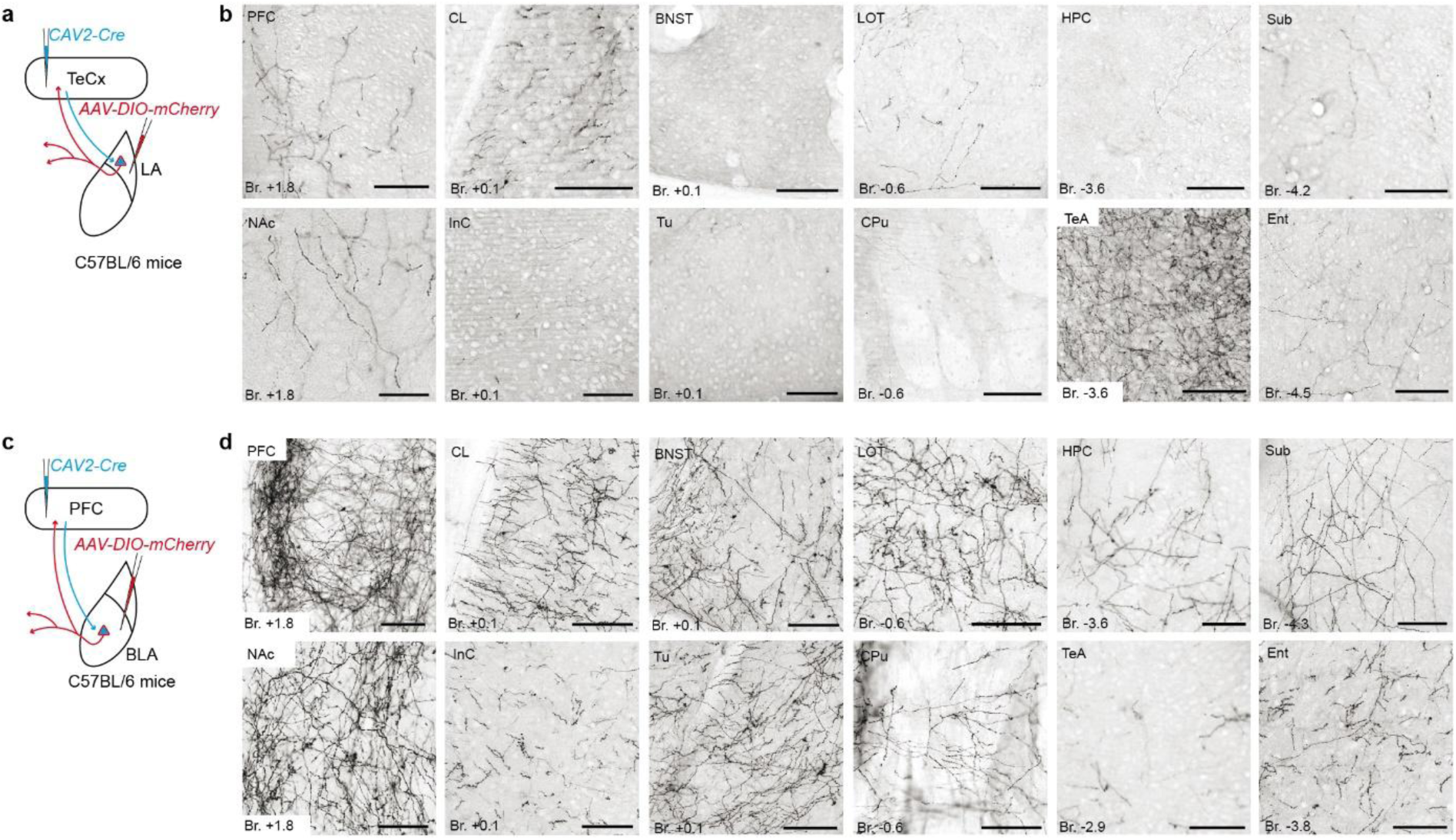
BLA, but not LA, innervates several extra-amygdala limbic regions (same trials as in Fig. 6) **a**,**c**, Experimental design for CAV2-Cre-mediated viral tracing of LA (a) and BLA neurons (c). **b**,**d**, High magnification brightfield images showing the distribution of mCherry-labelled LA (b) and BLA axons (d) throughout the forebrain. Scale bars, 100 µm. For abbreviations see *Supplementary Tables*.

**Extended data figure 18:**
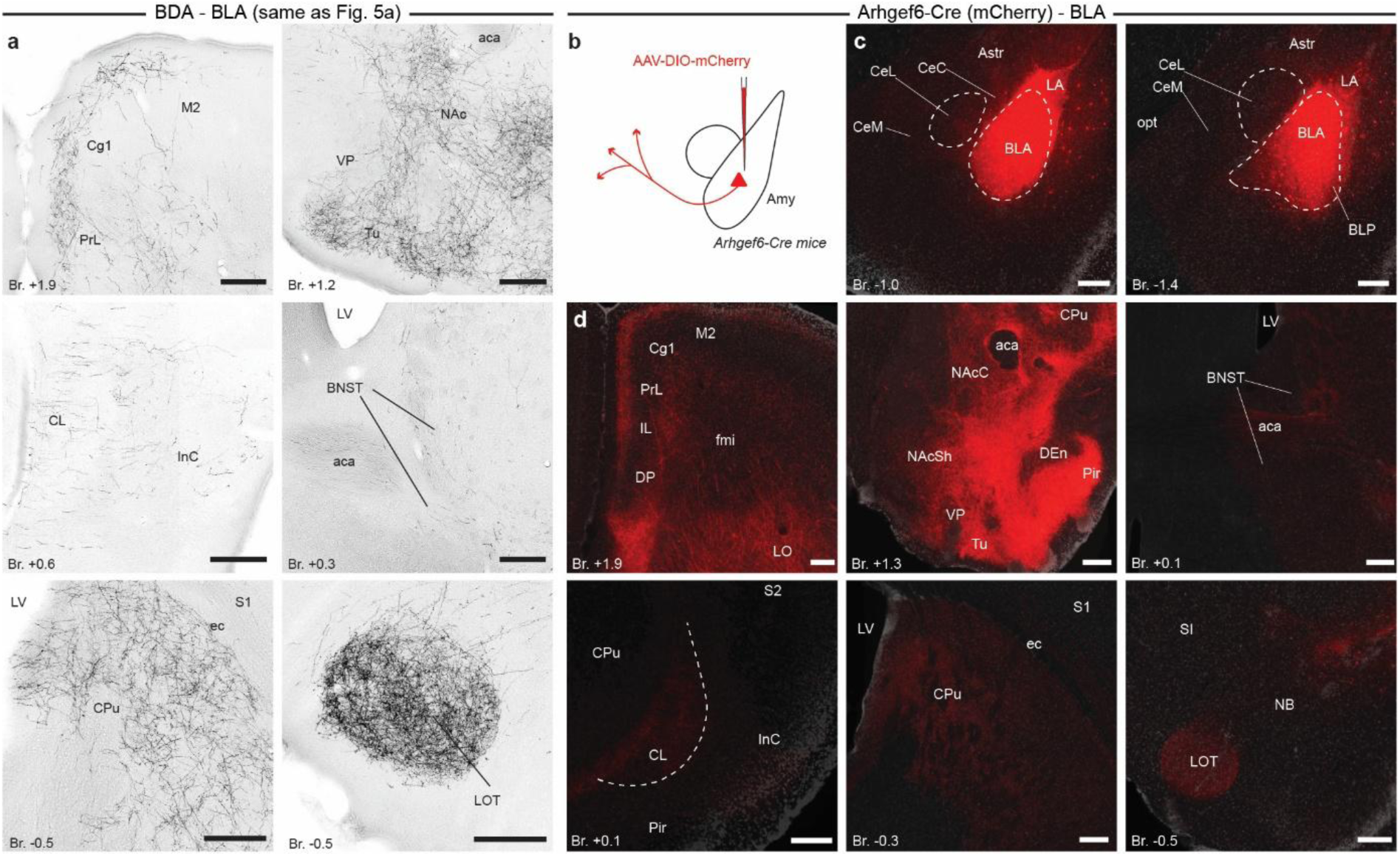
BLA innervates a wide range of limbic structures in the forebrain (related to Fig. 6) **a**, Brightfield light microscopic images obtained from the same trial as in Fig. 5a showing the distribution of BDA-labelled BLA axons in various limbic regions in the forebrain. Scale bars, 200 µm. **b,** Experimental design for viral tracing of BLA neurons in *Arhgef6*-Cre mice. **c**, mCherry-labelled neurons and their axons (red) in the amygdala. **d,** Distribution of viral-labelled BLA axons (red) throughout the forebrain. Scale bars, 200 µm. For abbreviations see *Supplementary Tables*.

## Notes

### Competing Interest Statement

The authors have declared no competing interest.

## References

1. LeDoux, J. E. Coming to terms with fear. Proceedings of the National Academy of Sciences 111, 2871–2878 (2014).

2. Fanselow, M. S. & Pennington, Z. T. A return to the psychiatric dark ages with a two-system framework for fear. Behaviour Research and Therapy 100, 24–29 (2018).

3. Zych, A. D. & Gogolla, N. Expressions of emotions across species. Curr. Opin. Neurobiol. 68, 57–66 (2021).

4. Davis, M. Neurobiology of fear responses: the role of the amygdala. J. Neuropsychiatry Clin. Neurosci. 9, 382–402 (1997).

5. LeDoux, J. The amygdala. Current Biology 17, R868–R874 (2007).

6. Janak, P. H. & Tye, K. M. From circuits to behaviour in the amygdala. Nature vol. 517 284–292 Preprint at 10.1038/nature14188 (2015).

7. Romanski, L. M. Gateway to the study of the amygdala and emotion. Cerebral Cortex 35, 3–4 (2025).

8. Lee, J. H., Lee, S. & Kim, J.-H. Amygdala Circuits for Fear Memory: A Key Role for Dopamine Regulation. The Neuroscientist 23, 542–553 (2017).

9. Kim, J. J. & Jung, M. W. Fear paradigms: The times they are a-changin’. Curr. Opin. Behav. Sci. 24, 38–43 (2018).

10. Liu, J., Totty, M. S., Bayer, H. & Maren, S. Integrating Aversive Memories in the Basolateral Amygdala. Biol. Psychiatry 10.1016/J.BIOPSYCH.2025.03.019 (2025) doi:10.1016/J.BIOPSYCH.2025.03.019.

11. Weinberger, N. M. The medial geniculate, not the amygdala, as the root of auditory fear conditioning. Hear. Res. 274, 61–74 (2011).

12. Do-Monte, F. H., Quiñones-Laracuente, K. & Quirk, G. J. A temporal shift in the circuits mediating retrieval of fear memory. Nature 519, 460–463 (2015).

13. Penzo, M. A. et al. The paraventricular thalamus controls a central amygdala fear circuit. Nature 519, 455–9 (2015).

14. Zhu, Y. et al. Dynamic salience processing in paraventricular thalamus gates associative learning. Science (1979). 362, 423–429 (2018).

15. Chen, M. & Bi, L. Optogenetic Long-Term Depression Induction in the PVT-CeL Circuitry Mediates Decreased Fear Memory. Mol. Neurobiol. 56, 4855–4865 (2019).

16. Barsy, B. et al. Associative and plastic thalamic signaling to the lateral amygdala controls fear behavior. Nat. Neurosci. 23, 625–637 (2020).

17. Chen, J. et al. The anterior paraventricular thalamus counteracts fear expression during retrieval through both amygdala and subiculum circuits. *Commun*. Biol. 8, 1635 (2025).

18. Taylor, J. A. et al. Single cell plasticity and population coding stability in auditory thalamus upon associative learning. Nat. Commun. 12, 2438 (2021).

19. Silva, B. A. et al. A thalamo-amygdalar circuit underlying the extinction of remote fear memories. Nat. Neurosci. 24, 964–974 (2021).

20. Cao, F. et al. Gustatory thalamic neurons mediate aversive behaviors. Nat. Commun. 16, 8517 (2025).

21. LeDoux, J., Farb, C. & Ruggiero, D. Topographic organization of neurons in the acoustic thalamus that project to the amygdala. The Journal of Neuroscience 10, 1043–1054 (1990).

22. Van der Werf, Y. D., Witter, M. P. & Groenewegen, H. J. The intralaminar and midline nuclei of the thalamus. Anatomical and functional evidence for participation in processes of arousal and awareness. Brain Res. Rev. 39, 107–140 (2002).

23. Mátyás, F. et al. A highly collateralized thalamic cell type with arousal-predicting activity serves as a key hub for graded state transitions in the forebrain. Nat. Neurosci. 21, 1551–1562 (2018).

24. Davis, M., Rainnie, D. & Cassell, M. Neurotransmission in the rat amygdala related to fear and anxiety. Trends Neurosci. 17, 208–214 (1994).

25. Orsini, C. A. & Maren, S. Neural and cellular mechanisms of fear and extinction memory formation. Neurosci. Biobehav. Rev. 36, 1773–1802 (2012).

26. Lee, S., Kim, S.-J., Kwon, O.-B., Lee, J. H. & Kim, J.-H. Inhibitory networks of the amygdala for emotional memory. Front. Neural Circuits 7, 58867 (2013).

27. Fenster, R. J., Lebois, L. A. M., Ressler, K. J. & Suh, J. Brain circuit dysfunction in post-traumatic stress disorder: from mouse to man. Nat. Rev. Neurosci. 10.1038/s41583-018-0039-7 (2018) doi:10.1038/s41583-018-0039-7.

28. Li, B. Central amygdala cells for learning and expressing aversive emotional memories. Curr. Opin. Behav. Sci. 26, 40–45 (2019).

29. Beyeler, A. et al. Divergent Routing of Positive and Negative Information from the Amygdala during Memory Retrieval. Neuron 90, 348–61 (2016).

30. Namburi, P. et al. A circuit mechanism for differentiating positive and negative associations. Nature 520, 675–678 (2015).

31. Beyeler, A. et al. Organization of Valence-Encoding and Projection-Defined Neurons in the Basolateral Amygdala. Cell Rep. 22, 905–918 (2018).

32. Kim, J., Pignatelli, M., Xu, S., Itohara, S. & Tonegawa, S. Antagonistic negative and positive neurons of the basolateral amygdala. Nat. Neurosci. 19, 1636–1646 (2016).

33. Kim, J., Zhang, X., Muralidhar, S., LeBlanc, S. A. & Tonegawa, S. Basolateral to Central Amygdala Neural Circuits for Appetitive Behaviors. Neuron 93, 1464–1479.e5 (2017).

34. Hintiryan, H. et al. Connectivity characterization of the mouse basolateral amygdalar complex. Nat. Commun. 12, 2859 (2021).

35. Romanski, L. M. & LeDoux, J. E. Information Cascade from Primary Auditory Cortex to the Amygdala: Corticocortical and Corticoamygdaloid Projections of Temporal Cortex in the Rat. Cerebral Cortex 3, 515–532 (1993).

36. Vertes, R. P. Differential projections of the infralimbic and prelimbic cortex in the rat. Synapse 51, 32–58 (2004).

37. LeDoux, J. E. Emotion Circuits in the Brain. Annu. Rev. Neurosci. 23, 155–184 (2000).

38. LeDoux, J. E. Semantics, Surplus Meaning, and the Science of Fear. Trends Cogn. Sci. 21, 303–306 (2017).

39. Mengxing, L., Lerma-Usabiaga, G., Clascá, F. & Paz-Alonso, P. M. High-Resolution Tractography Protocol to Investigate the Pathways between Human Mediodorsal Thalamic Nucleus and Prefrontal Cortex. The Journal of Neuroscience 43, 7780–7798 (2023).

40. Viena, T. D., Rasch, G. E., Silva, D. & Allen, T. A. Calretinin and calbindin architecture of the midline thalamus associated with prefrontal-hippocampal circuitry. Hippocampus 31, 770–789 (2021).

41. Fortin, M., Asselin, M. C., Gould, P. V. & Parent, A. Calretinin-immunoreactive neurons in the human thalamus. Neuroscience 84, 537–548 (1998).

42. Ledoux, J. E., Sakaguchi, A., Iwata, J. & Reis, D. J. Interruption of projections from the medial geniculate body to an archi-neostriatal field disrupts the classical conditioning of emotional responses to acoustic stimuli. Neuroscience 17, 615–627 (1986).

43. Gangarossa, G. et al. Contrasting patterns of ERK activation in the tail of the striatum in response to aversive and rewarding signals. J. Neurochem. 151, 204–226 (2019).

44. Mills, F. et al. Amygdalostriatal transition zone neurons encode sustained valence to direct conditioned behaviors. bioRxiv 2022.10.28.514263 Preprint at 10.1101/2022.10.28.514263 (2022).

45. Petersen, P. C., Siegle, J. H., Steinmetz, N. A., Mahallati, S. & Buzsáki, G. CellExplorer: A framework for visualizing and characterizing single neurons. Neuron 109, 3594–3608.e2 (2021).

46. Miller, L. M., Escabí, M. A., Read, H. L. & Schreiner, C. E. Functional convergence of response properties in the auditory thalamocortical system. Neuron 32, 151–160 (2001).

47. Sedigh-Sarvestani, M. et al. Intracellular, In Vivo, Dynamics of Thalamocortical Synapses in Visual Cortex. Journal of Neuroscience 37, 5250–5262 (2017).

48. Botta, P. et al. An Amygdala Circuit Mediates Experience-Dependent Momentary Arrests during Exploration. Cell 183, 605–619.e22 (2020).

49. Bordi, F. & LeDoux, J. E. Response properties of single units in areas of rat auditory thalamus that project to the amygdala. Exp. Brain Res. 98, 275–286 (1994).

50. Su, H.-S. & Bentivoglio, M. Thalamic midline cell populations projecting to the nucleus accumbens, amygdala, and hippocampus in the rat. J. Comp. Neurol. 297, 582–593 (1990).

51. Bordi, F. & Ledoux, J. Sensory Tuning beyond the Sensory System: An Initial Analysis of Auditory Response Properties of Neurons in the Lateral Amygdaloid Nucleus and Overlying Areas of the Striatum. The Journal of Neuroscience vol. 12 (1992).

52. Wang, T. et al. Paraventricular Thalamus Dynamically Modulates Aversive Memory via Tuning Prefrontal Inhibitory Circuitry. The Journal of Neuroscience 43, 3630 (2023).

53. Ren, S. et al. The paraventricular thalamus is a critical thalamic area for wakefulness. Science 362, 429–434 (2018).

54. Ma, C. et al. Sleep Regulation by Neurotensinergic Neurons in a Thalamo-Amygdala Circuit. Neuron 103, 323–334.e7 (2019).

55. Leppla, C. A. et al. Thalamus sends information about arousal but not valence to the amygdala. Psychopharmacology (Berl). 240, 477–499 (2023).

56. Wang, Y. et al. A common thalamic hub for general and defensive arousal control. Neuron 111, 3270–3287.e8 (2023).

57. Reeders, P. C., Rivera Núñez, M. V., Vertes, R. P., Mattfeld, A. T. & Allen, T. A. Identifying the midline thalamus in humans in vivo. Brain Struct. Funct. 228, 1835–1847 (2023).

58. Keifer, O. P., Gutman, D. A., Hecht, E. E., Keilholz, S. D. & Ressler, K. J. A comparative analysis of mouse and human medial geniculate nucleus connectivity: A DTI and anterograde tracing study. Neuroimage 105, 53–66 (2015).

59. Braunsdorf, M. et al. Does the temporal cortex make us human? A review of structural and functional diversity of the primate temporal lobe. Neurosci. Biobehav. Rev. 131, 400–410 (2021).

60. Homman-Ludiye, J. & Bourne, J. A. The medial pulvinar: function, origin and association with neurodevelopmental disorders. J. Anat. 235, 507–520 (2019).

61. Morán, M. A., Mufson, E. J. & Mesulam, M. -M. Neural inputs into the temporopolar cortex of the rhesus monkey. Journal of Comparative Neurology 256, 88–103 (1987).

62. Córdoba-Claros, M. A. et al. Projection Motifs and Wiring Logic of Medial Pulvinar Thalamocortical Axons in the Marmoset Monkey. Journal of Neuroscience 45, (2025).

63. Kark, S. M., Birnie, M. T., Baram, T. Z. & Yassa, M. A. Functional Connectivity of the Human Paraventricular Thalamic Nucleus: Insights From High Field Functional MRI. Front. Integr. Neurosci. 15, 662293 (2021).

64. Guedj, C. & Vuilleumier, P. Functional connectivity fingerprints of the human pulvinar: Decoding its role in cognition. Neuroimage 221, 117162 (2020).

65. Ressler, K. J. et al. Post-traumatic stress disorder: clinical and translational neuroscience from cells to circuits. Nature Reviews Neurology 2022 18:5 18, 273–288 (2022).

66. Calhoon, G. G. & Tye, K. M. Resolving the neural circuits of anxiety. Nat. Neurosci. 18, 1394–1404 (2015).

67. Jolkkonen, E., Pikkarainen, M., Kemppainen, S. & Pitkänen, A. Interconnectivity between the amygdaloid complex and the amygdalostriatal transition area: a PHA-L study in rat. J. Comp. Neurol. 431, 39–58 (2001).

68. Shammah-Lagnado, S. J., Alheid, G. F. & Heimer, L. Afferent connections of the interstitial nucleus of the posterior limb of the anterior commissure and adjacent amygdalostriatal transition area in the rat. Neuroscience 94, 1097–1123 (1999).

69. Wang, C., Wilson, W. A. & Moore, S. D. Role of NMDA, Non-NMDA, and GABA Receptors in Signal Propagation in the Amygdala Formation. J. Neurophysiol. 86, 1422–1429 (2001).

70. Wang, C., Kang-Park, M.-H., Wilson, W. A. & Moore, S. D. Properties of the Pathways From the Lateral Amygdal Nucleus to Basolateral Nucleus and Amygdalostriatal Transition Area. J. Neurophysiol. 87, 2593–2601 (2002).

71. Krettek, J. E. & Price, J. L. Amygdaloid projections to subcortical structures within the basal forebrain and brainstem in the rat and cat. Journal of Comparative Neurology 178, 225–253 (1978).

72. Kelley, A. E., Domesick, V. B. & Nauta, W. J. H. The amygdalostriatal projection in the rat—an anatomical study by anterograde and retrograde tracing methods. Neuroscience 7, 615–630 (1982).

73. Kita, H. & Kitai, S. T. Amygdaloid projections to the frontal cortex and the striatum in the rat. J. Comp. Neurol. 298, 40–49 (1990).

74. McDonald, A. J. Topographical organization of amygdaloid projections to the caudatoputamen, nucleus accumbens, and related striatal-like areas of the rat brain. Neuroscience 44, 15–33 (1991).

75. Stefanacci, L. et al. Projections from the lateral nucleus to the basal nucleus of the amygdala: A light and electron microscopic PHA-L study in the rat. Journal of Comparative Neurology 323, 586–601 (1992).

76. Pitkänen, A. et al. Intrinsic connections of the rat amygdaloid complex: Projections originating in the lateral nucleus. Journal of Comparative Neurology 356, 288–310 (1995).

77. Bourgeais, L., Gauriau, C. & Bernard, J. Projections from the nociceptive area of the central nucleus of the amygdala to the forebrain: a PHA-L study in the rat. European Journal of Neuroscience 14, 229–255 (2001).

78. De Armentia, M. L. & Sah, P. Firing properties and connectivity of neurons in the rat lateral central nucleus of the amygdala. J. Neurophysiol. 92, 1285–1294 (2004).

79. Killcross, S., Robbins, T. W. & Everitt, B. J. Different types of fear-conditioned behaviour mediated by separate nuclei within amygdala. Nature 388, 377–380 (1997).

80. Kishi, T., Tsumori, T., Yokota, S. & Yasui, Y. Topographical projection from the hippocampal formation to the amygdala: A combined anterograde and retrograde tracing study in the rat. J. Comp. Neurol. 496, 349–368 (2006).

81. Adhikari, A. et al. Basomedial amygdala mediates top-down control of anxiety and fear. Nature 527, 179–185 (2015).

82. Suh, J. et al. Projection-specific roles of basolateral amygdala Thy1 neurons in alcohol-induced place preference. Molecular Psychiatry 2025 1–11 (2025) doi:10.1038/s41380-025-03184-w.

83. Tian, Z. et al. The interhemispheric amygdala-accumbens circuit encodes negative valence in mice. Science (1979). 386, eadp7520 (2024).

84. Ren, J. et al. Single-cell transcriptomes and whole-brain projections of serotonin neurons in the mouse dorsal and median raphe nuclei. Elife 8, (2019).

85. Zingg, B., Dong, H. W., Tao, H. W. & Zhang, L. I. Input-output Organization of the Mouse Claustrum. J. Comp. Neurol. 526, 2428 (2018).

86. Aransay, A., Rodríguez-López, C., García-Amado, M., Clascá, F. & Prensa, L. Long-range projection neurons of the mouse ventral tegmental area: A single-cell axon tracing analysis. Front. Neuroanat. 9, 139323 (2015).

87. Sengupta, A. et al. Basolateral Amygdala Neurons Maintain Aversive Emotional Salience. The Journal of Neuroscience 38, 3001–3012 (2018).

88. Tang, W., Kochubey, O., Kintscher, M. & Schneggenburger, R. A VTA to basal amygdala dopamine projection contributes to signal salient somatosensory events during fear learning. The Journal of Neuroscience JN-RM-1796-19 (2020) doi:10.1523/JNEUROSCI.1796-19.2020.

89. Li, H. et al. Neurotensin orchestrates valence assignment in the amygdala. Nature 2022 608:7923 608, 586–592 (2022).

90. Burgos-Robles, A. et al. Amygdala inputs to prefrontal cortex guide behavior amid conflicting cues of reward and punishment. Nat. Neurosci. 20, 824–835 (2017).

91. McNally, G. P. Motivational competition and the paraventricular thalamus. Neurosci. Biobehav. Rev. 125, 193–207 (2021).

92. Millan, E. Z., Ong, Z. & McNally, G. P. Paraventricular thalamus: Gateway to feeding, appetitive motivation, and drug addiction. in Progress in Brain Research vol. 235 113–137 (Elsevier, 2017).

93. Paxinos, G. & Franklin, K. B. J. The Mouse Brain in Stereotaxic Coordinates*, 5th Edition*. (Academic Press, 2019).

94. Allen Reference Atlas – Mouse Brain. Allen Reference Atlas – Mouse Brain. https://atlas.brain-map.org/.

95. Herry, C. et al. Switching on and off fear by distinct neuronal circuits. Nature 454, 600–606 (2008).

96. Polepalli, J. S., Gooch, H. & Sah, P. Diversity of interneurons in the lateral and basal amygdala. NPJ Sci. Learn. 5, 10 (2020).

97. Bukalo, O. et al. Prefrontal inputs to the amygdala instruct fear extinction memory formation. Sci. Adv. 1, e1500251–e1500251 (2015).

98. Anglada-Figueroa, D. & Quirk, G. J. Lesions of the Basal Amygdala Block Expression of Conditioned Fear But Not Extinction. The Journal of Neuroscience 25, 9680–9685 (2005).

99. Manassero, E., Renna, A., Milano, L. & Sacchetti, B. Lateral and Basal Amygdala Account for Opposite Behavioral Responses during the Long-Term Expression of Fearful Memories. Sci. Rep. 8, 518 (2018).

100. Kim, W. Bin & Cho, J. H. Encoding of contextual fear memory in hippocampal–amygdala circuit. Nat. Commun. 11, 1–22 (2020).

101. Tye, K. M. et al. Amygdala circuitry mediating reversible and bidirectional control of anxiety. Nature 471, 358–362 (2011).

102. Terburg, D. et al. The Basolateral Amygdala Is Essential for Rapid Escape: A Human and Rodent Study. Cell 175, 723–735.e16 (2018).

103. Asede, D., Doddapaneni, D. & Bolton, M. M. Amygdala Intercalated Cells: Gate Keepers and Conveyors of Internal State to the Circuits of Emotion. The Journal of Neuroscience 42, 9098–9109 (2022).

104. Chaaya, N. et al. Localization of Contextual and Context Removed Auditory Fear Memory within the Basolateral Amygdala Complex. Neuroscience 398, 231–251 (2019).

105. Gründemann, J. & Lüthi, A. Ensemble coding in amygdala circuits for associative learning. Curr. Opin. Neurobiol. 35, 200–206 (2015).

106. Rajbhandari, A. K. et al. A Basomedial Amygdala to Intercalated Cells Microcircuit Expressing PACAP and Its Receptor PAC1 Regulates Contextual Fear. The Journal of Neuroscience 41, 3446–3461 (2021).

107. McDonald, A. J., Shammah-Lagnado, S. J., Shi, C. & Davis, M. Cortical afferents to the extended amygdala. Ann. N. Y. Acad. Sci. 877, 309–38 (1999).

108. Bordi, F., LeDoux, J., Clugnet, M. C. & Pavlides, C. Single-unit activity in the lateral nucleus of the amygdala and overlying areas of the striatum in freely behaving rats: Rates, discharge patterns, and responses to acoustic stimuli. Behavioral Neuroscience 107, 757–769 (1993).

109. Turner, B. H. & Zimmer, J. The architecture and some of the interconnections of the rat’s amygdala and lateral periallocortex. J. Comp. Neurol. 227, 540–557 (1984).

110. LeDoux, J. E., Farb, C. R. & Romanski, L. M. Overlapping projections to the amygdala and striatum from auditory processing areas of the thalamus and cortex. Neurosci. Lett. 134, 139–144 (1991).

111. Mascagni, F., McDonald, A. J. & Coleman, J. R. Corticoamygdaloid and corticocortical projections of the rat temporal cortex: APhaseolus vulgaris leucoagglutinin study. Neuroscience 57, 697–715 (1993).

112. Moga, M. M., Weis, R. P. & Moore, R. Y. Efferent projections of the paraventricular thalamic nucleus in the rat. J. Comp. Neurol. 359, 221–238 (1995).

113. Shi, C.-J. & Cassell, M. D. Cortical, thalamic, and amygdaloid connections of the anterior and posterior insular cortices. J. Comp. Neurol. 399, 440–468 (1998).

114. Bernard, J. -F, Alden, M. & Besson, J. -M. The organization of the efferent projections from the pontine parabrachial area to the amygdaloid complex: A phaseolus vulgaris leucoagglutinin (PHA-L) study in the rat. Journal of Comparative Neurology 329, 201–229 (1993).

115. Jasnow, A. M. et al. Thy1-Expressing Neurons in the Basolateral Amygdala May Mediate Fear Inhibition. Journal of Neuroscience **June**, 10396–10404 (2013).

116. Babiczky, Á. & Matyas, F. Molecular characteristics and laminar distribution of prefrontal neurons projecting to the mesolimbic system. Elife 11, (2022).

117. Katona, I. et al. Distribution of CB1 Cannabinoid Receptors in the Amygdala and their Role in the Control of GABAergic Transmission. The Journal of Neuroscience 21, 9506–9518 (2001).

118. Haubensak, W. et al. Genetic dissection of an amygdala microcircuit that gates conditioned fear. Nature 468, 270–276 (2010).

119. Penzo, M. A., Robert, V. & Li, B. Fear Conditioning Potentiates Synaptic Transmission onto Long-Range Projection Neurons in the Lateral Subdivision of Central Amygdala. The Journal of Neuroscience 34, 2432–2437 (2014).

120. Han, S., Soleiman, M. T., Soden, M. E., Zweifel, L. S. & Palmiter, R. D. Elucidating an Affective Pain Circuit that Creates a Threat Memory. Cell 162, 363–374 (2015).

121. Hagihara, K. M. et al. Intercalated amygdala clusters orchestrate a switch in fear state. Nature 594, 403–407 (2021).

122. Barabás, B., Reéb, Z., Papp, O. I. & Hájos, N. Functionally linked amygdala and prefrontal cortical regions are innervated by both single and double projecting cholinergic neurons. Front. Cell. Neurosci. 18, (2024).

123. Lerma-Usabiaga, G., Liu, M., Paz-Alonso, P. M. & Wandell, B. A. Reproducible Tract Profiles 2 (RTP2) suite, from diffusion MRI acquisition to clinical practice and research. Scientific Reports 2023 13:1 13, 6010- (2023).

124. Iglesias, J. E. et al. A probabilistic atlas of the human thalamic nuclei combining ex vivo MRI and histology. Neuroimage 183, 314–326 (2018).

125. Glasser, M. F. et al. A multi-modal parcellation of human cerebral cortex. Nature 2016 536:7615 536, 171–178 (2016).

126. Tournier, J. D. et al. MRtrix3: A fast, flexible and open software framework for medical image processing and visualisation. Neuroimage 202, 116137 (2019).

127. Veraart, J. et al. Denoising of diffusion MRI using random matrix theory. Neuroimage 142, 394–406 (2016).

128. Kellner, E., Dhital, B., Kiselev, V. G. & Reisert, M. Gibbs-ringing artifact removal based on local subvoxel-shifts. Magn. Reson. Med. 76, 1574–1581 (2016).

129. Smith, S. M. et al. Advances in functional and structural MR image analysis and implementation as FSL. Neuroimage 23, S208–S219 (2004).

130. Jeurissen, B., Tournier, J. D., Dhollander, T., Connelly, A. & Sijbers, J. Multi-tissue constrained spherical deconvolution for improved analysis of multi-shell diffusion MRI data. Neuroimage 103, 411–426 (2014).

131. Tournier, J.-D., Calamante, F. & Connelly, A. Improved Probabilistic Streamlines Tractography by 2nd Order Integration Over Fibre Orientation Distributions. in Proceedings of the International Society for Magnetic Resonance in Medicine 1670 (Stockholm, 2010).

132. Pachitariu, M., Sridhar, S., Pennington, J. & Stringer, C. Spike sorting with Kilosort4. Nature Methods 2024 21:5 21, 914–921 (2024).

133. Henze, D. A. et al. Intracellular features predicted by extracellular recordings in the hippocampus in vivo. J. Neurophysiol. 84, 390–400 (2000).

